# Reducing the efforts to create reproducible analysis code with FieldTrip

**DOI:** 10.1101/2021.02.05.429886

**Authors:** Mats W.J. van Es, Eelke Spaak, Jan-Mathijs Schoffelen, Robert Oostenveld

## Abstract

The analysis of EEG and MEG data typically requires a lengthy and complicated sequence of analysis steps, often requiring large amounts of computations, which are ideally represented in analysis scripts. These scripts are often written by researchers without formal training in computer science, resulting in the quality and readability of these analysis scripts to be highly dependent on individual coding expertise and style. Even though the computational outcomes and interpretation of the results can be correct, the inconsistent style and quality of analysis scripts make reviewing the details of the analysis difficult for other researchers that are either involved in the study or not, and the quality of the scripts might compromise the reproducibility of obtained results. This paper describes the design and implementation of a strategy that allows complete reproduction of MATLAB-based scripts with little extra efforts on behalf of the user, which we have implemented as part of the FieldTrip toolbox. Starting from the researchers’ idiosyncratic pipeline scripts, this new functionality allows researchers to automatically create and publish analysis pipeline scripts in a standardized format, along with all relevant intermediate data. We demonstrate the functionality and validate its effectiveness by applying it to the analysis of a recently published MEG study.

## Introduction

Unsound scientific practices have led to a replication crisis in psychological science in recent years (1,2), and it is unlikely that cognitive neuroscience is an exception (3–5). Initiatives to combat this crisis are taking root (4,6,7), targeted at increasing robustness of results, publishing of null results, and greater methodological transparency. This has resulted in publications with recommendations for best practices (8,9), but these have not yet been universally embraced. The increased sophistication of experimental designs and analysis methods also results in data analysis getting so complex that the methods sections of manuscripts in most journals is too short to represent the analysis in sufficient detail, thus hampering transparency. Therefore, researchers are increasingly encouraged to share their data and analysis pipelines along with their published results (6).

### Scientific analysis represented as scripts and pipelines

The processing and analysis of EEG and MEG data typically involves a sequence of steps, often requiring large amounts of computations. Each of these steps is based on input data, and produces output data, hence these analyses can be conceptualized as pipelines through which data “flows”, where each stage modifies the data somehow. Overall, the input to the analysis pipeline comprises raw data, and the output consists of interpretable results. The steps in an analysis pipeline are typically represented as code in a human-readable programming language such as MATLAB, Python or Julia (*source code*), in files called *scripts*. The quality, readability, and generalizability of the scripts, which are written by individual researchers, is highly dependent on individual coding style and expertise. Since the reproducibility of the pipeline depends on the quality of the analysis scripts, variability in the quality might compromise the reproducibility of obtained results. Furthermore, since scripts might be difficult to read, it can be problematic to find (and learn from) the details of the analytical procedures applied in previous studies. In practice we also see that researchers set high standards for themselves and therefore are hesitant to openly share their own analysis code, because the code is not as clean and well-documented as they would like.

### Pipeline systems

A number of strategies have been proposed to enhance the reproducibility of analysis pipelines and scientific results. One option to improve reproducibility and efficiency through reuse of code is through automation using pipeline systems (e.g. Taverna, Galaxy, LONI, PSOM, Nipype, Brainlife; (10–15) or batch scripts (e.g. SPM’s matlabbatch (16)). Generally, these provide the researcher with tools to construct an analysis pipeline, manage the execution of the steps in the pipeline and, to a varying degree, handle data. The pipeline system manages the execution of code and automatically passes the data from one analysis step to the next, even when these are implemented in different analysis software. Besides providing a better visual and conceptual overview of elaborate pipelines and improving the efficiency of the researcher’s workflow, pipeline systems improve reproducibility by providing an explicitly specified sequence of analysis steps, combined with detailed output logs.

Some drawbacks of pipeline systems are that they require the researcher to learn how the pipeline software works on top of leaning the analysis itself, that the execution requires extra software to be installed, or that it requires moving the execution from a local computer to an online (cluster or cloud-based) system, and that the flexibility of pipeline systems is limited. Some data analysis strategies can not easily be translated to fully automatable pipelines, because manual intervention and interactive examination of data or intermediate results is required. There are however also platforms (e.g. brainlife (14)) that enable interaction, while maintaining the advantages of a pipeline system. Furthermore, the researcher is limited to using the tools that are already available in a specific pipeline system, or is required to extend the pipeline system themselves by writing wrappers between the pipeline system and the analysis tools of choice. Moreover, the pipeline system only improves reproducibility perfectly when the dependencies of the system, e.g. specific versions of external toolboxes, template data, and external (web-based) services, are well defined. If the pipeline depends on, or allows the execution of, custom analysis steps (thereby providing maximum analytical freedom), the source code of those custom analysis steps is also required.

### Version control systems for scripts

If an analysis pipeline depends on custom scripts, it is vital for reproducibility to document the version of the code that produced the result. Version Control Systems (VCS) can facilitate this, by providing tools to track and control changes made to source code (17–20). Especially the Git version control system is increasingly being used in science (21), which is in part due to GitHub providing a popular online web platform that gives a clear graphical presentation of projects and code, and facilitates collaboration (“social coding”) and dissemination. In a VCS, a complete history of the incremental changes to the source code is saved, and each revision is given a unique identifier. This enables code developers to compare versions and retrace errors, but also facilitates multiple developers to contribute to the same project and merge contributions (21). When using a VCS for scientific workflows, these can easily be shared and published. This has the potential to increase the reproducibility of a scientific project, but only when a number of conditions are met.

First, the researcher has to use the VCS tool actively on their own code. Working with VCSs requires training (20), and even if a researcher is well-trained, they have to commit a new version of the source code after each significant change to the code, ideally including a short textual description of the changes that were made. This style of working involves extra time investment, and is easily given a lower priority in the busy day-to-day work of a researcher and therefore skipped.

Second, for the pipeline to be reproducible for outside parties, the researcher has to be willing to share their analysis workflow. Researchers might be hesitant to do so if they feel insecure about their coding style and the quality of their code. During their academic training, researchers learn how to present results and write papers, and receive positive reinforcement on these skills; the development of computing and coding skills are often not part of the formal training programs for young neuroscientists, or are not as well established (22,23). Low quality code might reflect badly on the quality of their scientific work, and hence researchers might feel obliged to invest extra time to clean up and document their code solely for sharing purposes. In our personal experience, it is not uncommon to hear colleagues and coworkers that they will share the code in the future, after it has been cleaned up and documented, but that in the end the code is never shared. Training researchers in pro-actively writing clean code and documenting it definitely is of help here, and we encourage researchers to study examples and use experts’ tips and tricks (e.g. 22, 23). However, given time constraints, prioritization of other aspects related to the research project, or simply the fact that the researcher does not know of the existence of such guidelines, it can happen that the researcher decides not to share the scripts at all. In a trend we view as positive, journals and publishers are increasingly demanding transparency when researchers wish to publish their results, and one of these requirements can be to provide access to data and analysis scripts (26).

Last, the workflow that is tracked in a VCS needs to be executable by other researchers on other computers. This can be challenging, especially if the researcher’s scripts depend on external code and toolboxes that by themselves are not tracked by the VCS. Similarly, analysis scripts typically depend on a particular organization of the data over directories and sub-directories, which is unknown to outsiders. Tools to document code dependencies are becoming more widely adopted, such as Conda, and Python’s virtualenv, modulefinder, and “pip freeze”. Furthermore, the problem of ill-specified data organization can (partly) be overcome by recently developed data organization standards such as the Brain Imaging Data Structure (BIDS, (18,27,28). To summarize, sharing reproducible analysis pipelines can be challenging given that researchers may not be version controlling their own code so well, may not be making their own code available, and because of issues created by unknown dependencies on untracked code and undocumented features of the data organization. The latter two issues might be overcome with platforms such as Code Ocean (29), which is an open access platform where users can develop their code and run their analysis in the same environment, but this still leaves the first issue unscathed.

### Literate programming

There are multiple styles of computer programming that lead to different degrees of reproducibility. The easiest way of programming is by using read-eval-print loops (REPLs), such as employed in the MATLAB (30) command window. These take single user inputs, evaluate them, and return the result to the user. Only using the command window or interpreter to execute REPLs is bad for reproducibility, since the details and sequence of analysis steps is not documented. One can improve upon this by saving the code in scripts. It is even better to include inline documentation (“comments”) in these scripts, describing the rationale of the code. The script then becomes a combination of a programming language targeted at a computer, and a documentation language targeted at peers, thereby improving the interpretability of the researcher’s code. This is known as literate programming (31) and it improves the transparency regarding how source code leads to results. Tools and packages designed for interactive literate programming are available for different scientific programming languages (e.g. MATLAB live scripts, Jupyter Notebook, R Markdown, knitr, and matlabweb (32–36)) and allow users to execute pieces of code and look at the results as a REPL, while simultaneously encouraging inline documentation, keeping a trace of all computational steps, and providing the ability to revisit and share interactive analyses. Online shared notebooks often make re-execution of the code possible without the need to install software packages or to download data. Combining REPL with literate programming benefits both the reproducibility of single analysis steps, and the transparency of the scientific process. However, it has been noted by users of Jupyter Notebooks that the interface becomes less manageable as the content grows, which drives users to clean up their notebooks and only keep the successful steps (37). Additionally, most users only annotate their code after writing it, solely when they want to share the code. Consequently, the longer, the more sophisticated, and the more interactive an analysis pipeline becomes, the less transparent it becomes. Finally, the integration of literate programming REPL code with VCSs is not always optimal.

### It all takes time

The improvement of reproducibility by the use of pipeline systems, VCSs, or by literate programming tools relies on the researcher using these tools properly and consistently. Moreover, it necessitates extra time investment on the researcher’s part in order to make their data and analysis scripts shareable. While we recommend the use of such tools, these are currently not (yet) widely adopted. The tools with the highest chance of adoption are usually the ones with the least friction, i.e. the least effort on the researchers’ part. Ideally, researchers should be able to transform their (often highly idiosyncratic) analysis scripts into a standard pipeline format automatically, allowing exact and transparent reproducibility.

We here present an implementation of such functionality in the FieldTrip toolbox (38), which is currently one of the most widely used toolboxes for MEG and EEG analysis. Using this new functionality, data analysis within the FieldTrip and MATLAB ecosystem can be made entirely reproducible and transparent with minimal additional effort by the researchers.

### Our solution

Our primary goal is to make analyses reproducible and to allow researchers to easily share details of their analysis pipeline, yet require minimal extra time investment or training of the researcher. We implemented this in the form of what we call the ‘*reproducescript*’ functionality. In short: the researcher adds one additional flag to the configuration options in each FieldTrip function in the pipeline, which results in the analysis pipeline and data dependencies to be exported to a standardized representation that resembles the format of the FieldTrip tutorials which the researcher will be familiar with. The generated scripts and corresponding data have minimal to no ambiguity. By standardizing the coding style used in scripts, researchers do not have to worry about the quality of their personal coding style. All the while, the analysis flexibility inherent to the FieldTrip toolbox remains, including interactive analysis steps. Finally, *reproducescript* enables the researcher to gradually build up and execute parts of their analysis using the approach they are used to, without the need to compile a complete pipeline at the start (e.g., preprocessing can be completed before the rest of the analysis pipeline is in place).

Specifically, upon the deployment of a processing pipeline (based on an individual script or collection of scripts), *reproducescript* saves all intermediate data from each analysis step in one place, according to a uniform file naming scheme and directory structure. Intermediate data include the input data to each individual analysis step, as well as the output each step generates. Additionally, reproducescript creates a standard-syntax, human-readable, executable, MATLAB script that solely relies on the FieldTrip toolbox, which, together with the intermediate data, can fully reproduce the entire analysis pipeline with one single command or mouse click. Researchers enable *reproducescript* with a single global configuration option, and can archive and opt to share the generated code, instead of (or in addition to) their custom-written code. In the remainder of this article, we will explain how this functionality is used, and demonstrate its ability using examples from published research in which the FieldTrip toolbox was used for analysis. Further limitations and considerations will also be discussed.

## Reproducescript

To explain the new *reproducescript* functionality, we will demonstrate its use it with a simple example pipeline for a single-subject analysis that comprises only a few analysis steps. For this we assume the reader already to be familiar with the structure of FieldTrip toolbox functions (see 33) and the way these are used during analysis. The example also demonstrates how it is employed by the researcher. Second, we demonstrate its application in a complete pipeline with preprocessing for multiple subjects, followed by a group analysis. The original idiosyncratic scripts that we selected for these first two examples are relatively clean and transparent, which means they are easily reproducible even without the new *reproducescript* functionality. Thus, they solely function as practical demonstrations. As an analysis pipeline becomes more complex, especially when it starts to contain various layers of custom functions over FieldTrip functions, the researcher’s original code can become more opaque. In such cases, the advantage of *reproducescript* to improve the readability and reproducibility becomes more apparent. As a final, third, example, we will therefore apply it to an already published analysis pipeline that contains such complexity. The analysis code and data used in these examples are publicly available in the Donders Repository (https://doi.org/10.34973/21pa-dg13).

### How it works

#### Example 1: single-subject analysis

##### Original analysis

To show how the *reproducescript* functionality works, we apply it to a script from the tutorial “Trigger-based trial selection” that is available on the FieldTrip website http://www.fieldtriptoolbox.org/tutorial/preprocessing/). The directory tree used in this example and the original source code are shown in Figure 1 and listing 1, respectively. The *reproducescript* functionality is initiated by the source code in listing 2, and when applied to the original source code (listing 1) it generates the files shown in figure 2 and the code shown in listing 3.

**Figure 1.**
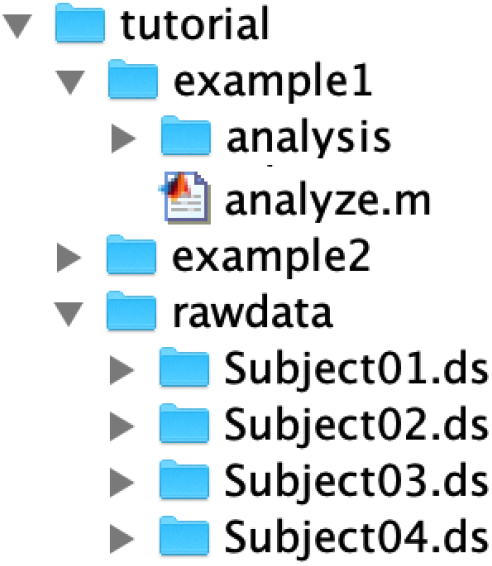
File directory tree. The *tutorial* folder contains subfolders for example 1 and 2, and the folder for the raw data used in both examples. The file *analyze.m* contains the analysis and saves the result in the *analysis* folder.

**Figure 2.**
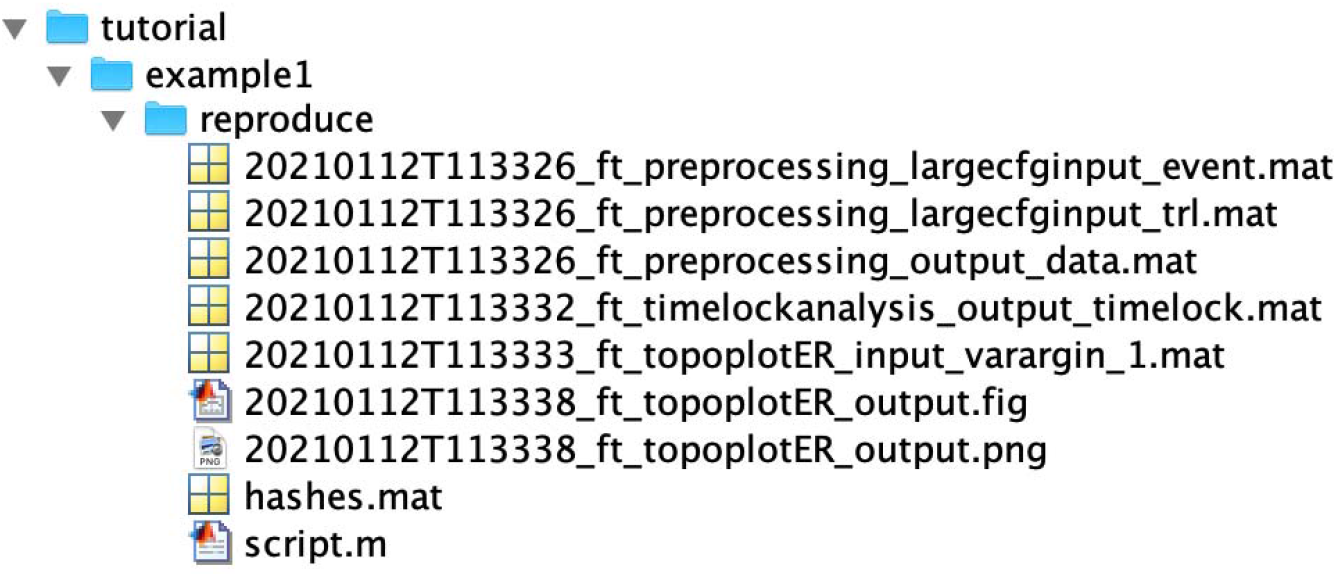
*reproducescript* creates the *reproduce* folder and its contents: input and output data files with unique file identifiers, a MATLAB script, and a *hashes* data file. See text for further explanation.

**Listing 1.**
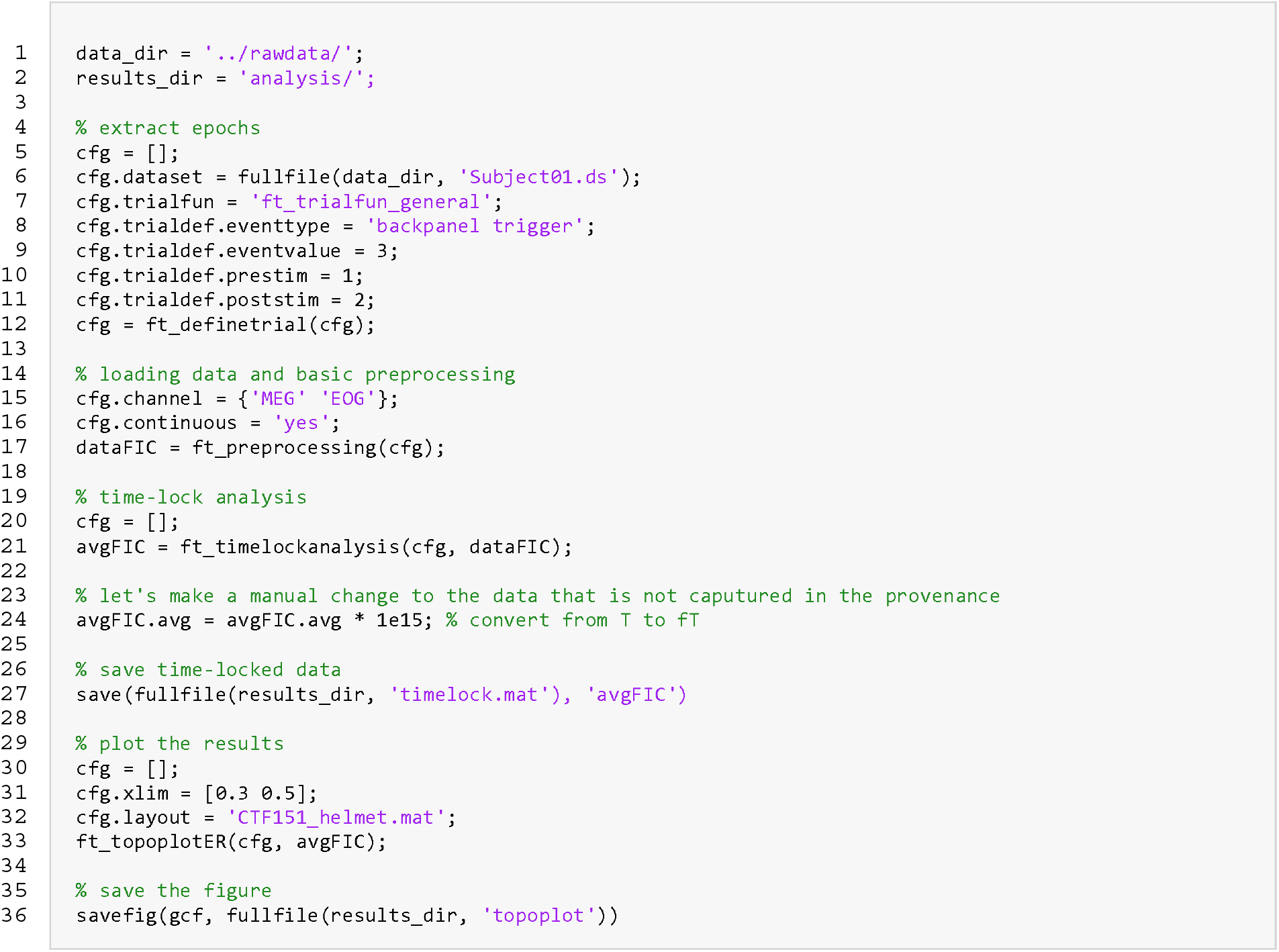
Example single-subject analysis from the FieldTrip tutorial. The script calls two FieldTrip functions for extracting epochs (*ft_definetrial*) and for reading in and preprocessing the data (*ft_preprocessing*). The data are averaged over trials (*ft_timelockanalysis*), manually transformed to femtotesla (fT), saved and visualized (*ft_topoplotER*).

A MATLAB analysis script that builds on the FieldTrip toolbox consists of a sequence of calls to FieldTrip functions, each of which perform a conceptual step of the analysis. The first input argument to such function is a configuration structure (*cfg*), which specifies the settings and parameters used by the function, and a data structure can be given as a subsequent input argument. The application of the function’s algorithms on the input data generates an output data structure. This can serve as input data to the next analysis step, or as the final result, in which case results can be visualized using a plotting function. Data structures are commonly represented in MATLAB memory, but can also be stored on disk in a *.mat file that is based on HDF5. In this example, we use the first analysis steps that are used in a typical pipeline. First, extraction of epochs of interest is accomplished using *ft_definetrial*. Its output can be used by *ft_preprocessing* to read the data from disk and do basic preprocessing. Finally, *ft_timelockanalysis* computes the average over trials, and *ft_topoplotER* plots the results.

Both the input and output of FieldTrip functions can be a data structure and/or a *cfg* structure. The *ft_definetrial* call only requires a configuration structure to be specified, including the directory of the raw data. Its output, too, is only a *cfg-structure*, which among others now contains a *trl*-field with the relevant information for epoching the data. The output from *ft_preprocessing* and *ft_timelockanalysis* are data structures, executed according to the configuration options specified in *cfg*. The only output from *ft_topoplotER* is a figure. Before calling *ft_topoplotER*, we changed the units from T to fT. This is usually not done, but in this instance it serves as an example for how *reproducescript* handles analysis steps that were performed outside the FieldTrip ecosystem (i.e., arbitrary code).

##### Initialization of reproducescript

**Listing 2.**
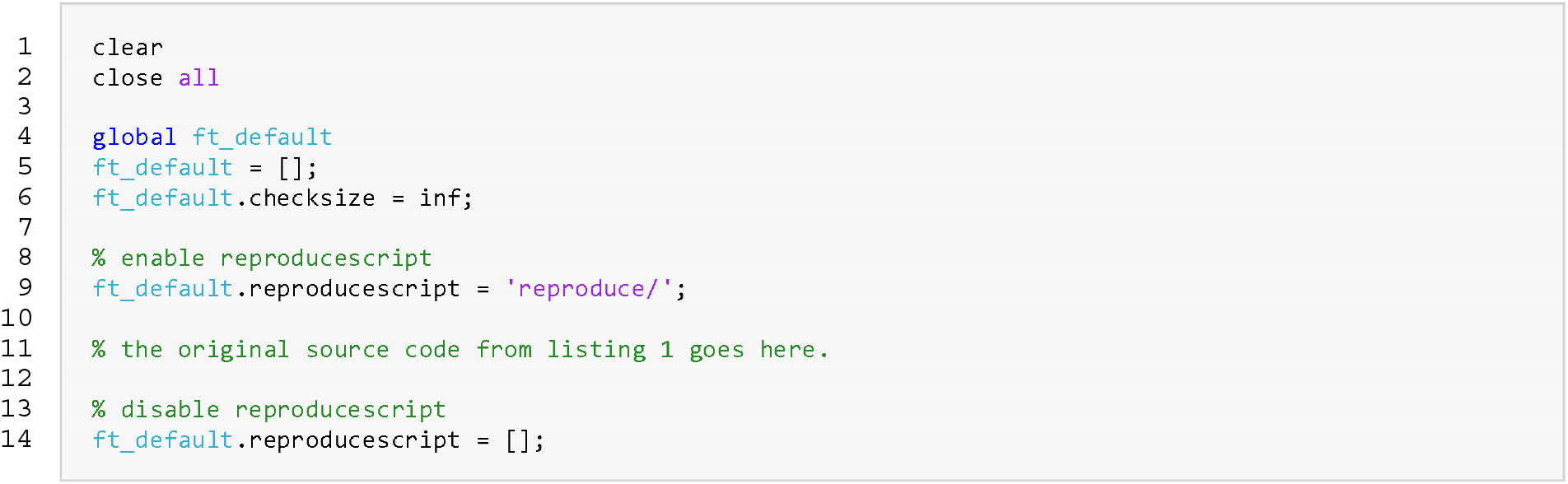
*reproducescript* is initialized at the top of the analysis script, by specifying the directory of the *reproducescript* folder in *ft_default.reproducescript*.

The functionality for reproducibility of analysis pipelines in the FieldTrip ecosystem is enabled at the top of a script (listing 2). The user specifies the directory to which the standard script and intermediate data are written in the reproducescript field of the global *ft_default* variable. *ft_default* is the structure in which global configuration defaults are stored; it is used throughout all FieldTrip functions and global options are at the start of the function merged with the user-supplied options in the *cfg* structure specific to the function. Note that we are additionally specifying ‘*ft_default.checksize = inf*’, which instructs FieldTrip to never remove (large) fields from any cfg-structure, thus ensuring perfect reproducibility. We recommend enabling this additional option whenever *reproducescript* is used.

##### Reproduced analysis

The *reproducescript* option is enabled by specifying an output directory in each functions *cfg* structure or in the global *ft_default* variable if we want it to apply to all functions. FieldTrip functions that are subsequently called will ensure that the output directory exists, and will store the relevant files in this directory (figure 2). *reproducescript* traces the steps to each FieldTrip function call, and recreates human-readable REPL code from scratch (script.m, listing 3). At the same time, the input data to a function and the output data it generates are copied and given a unique identifier (i.e. filenames). Pointers to these identifiers end up in the standardized script (listing 3), as *cfg.inputfile* and *cfg.outputfile*. This means that no input or output data structures as they normally appear in the MATLAB workspace appear in the standardized script; these are all handled using data on disk corresponding with *cfg.inputfile* and *cfg.outputfile*. In the case of *ft_definetrial*, these are absent, because the input and output of *ft_definetrial* are only *cfg* structures, not data structures. Similarly, if the function’s output is a figure (e.g. in *ft_topoplotER*) the figure is also directly saved to disk, in .png (bitmap) and .fig (MATLAB figure) formats.

Note that the fields from the *cfg* input to *ft_definetrial* are repeated as input to *ft_preprocessing* because the configuration in the original script was not emptied (listing 1, line 15). There are also additional fields created by *ft_definetrial*. If these fields exceed a certain printed size, which would make them unwieldy to include inline in a script (e.g. *cfg.trl*, which normally consists of a Ntrials*3 matrix specifying the relevant sections of the data on disk), these too are saved on disk instead of being printed in the standardized script. One last thing that should stand out is the comment in listing 3, line 47: “a new input variable is entering the pipeline here: …”. This points to the mat-file subsequently specified in *cfg.inputfile* to *ft_topoplotER*. The data structure in this file was not originally created by a FieldTrip function but comes from another source: in this case it consists of the data in which originates from the T to fT unit conversion step (listing 1, line 24). Thus, this comment puts an emphasis on the fact that a data structure with unknown provenance enters the pipeline. All analysis steps that do not use FieldTrip functions will create such comments and save the data structure. Importantly, the pipeline thus remains reproducible without relying on external code (see Discussion).

**Listing 3.**
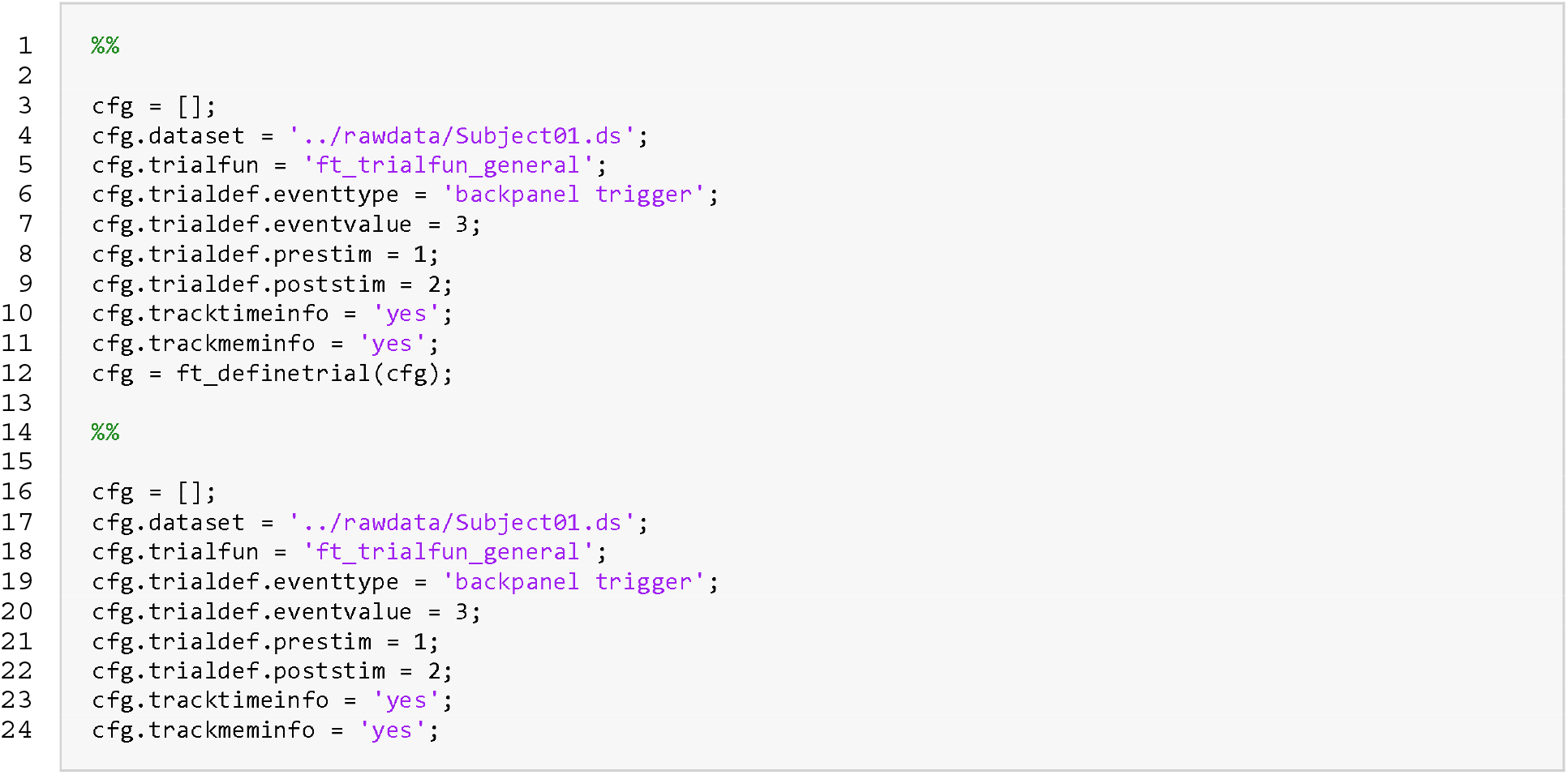
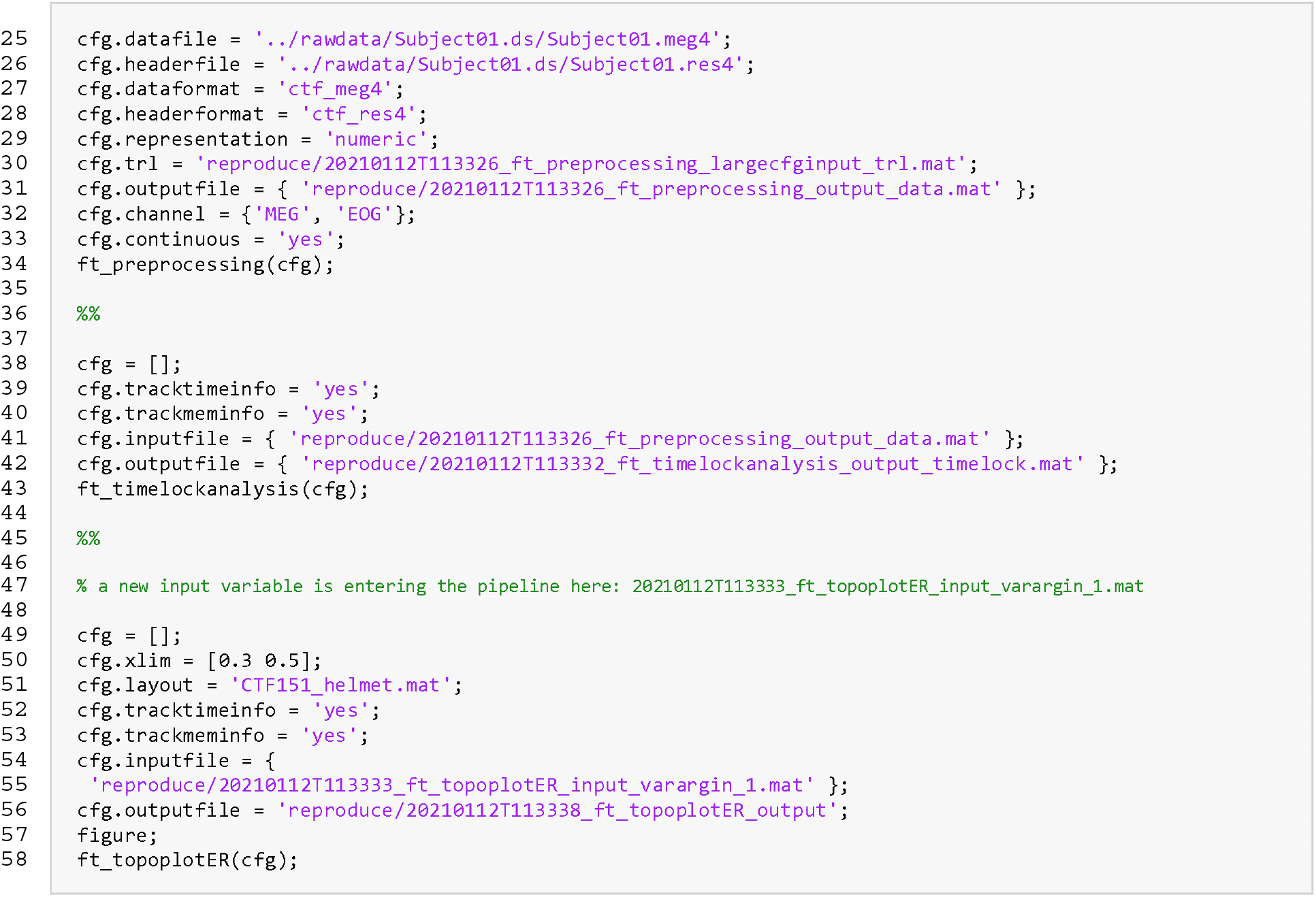
Example reproducescript output. This script is generated by *reproducescript* when listing 1 and 2 are combined and executed.

Finally, the *reproduce* folder contains a file named *hashes.mat*. This is a file containing MD5 hashes for bookkeeping all input and output files. It allows reproducescript to match the output files of any one step to the input files of any subsequent step. For example, the output from *ft_preprocessing* is used as input to *ft_timelockanalysis*, which means that the data structure only needs to be stored once and “…_ft_timelockanalysis_*input*_timelock.mat” does not have to be additionally saved to disk. If the output data from one function and the input data to the next function are slightly different, they are both saved under different file names. This happens when the researcher modified the data using custom code (as in the example when converting channel units). The *hashes.mat* file furthermore allows any researcher to check the integrity of all the intermediate and final result files of the pipeline.

#### Example 2: group analysis

The first example contained only a few analysis steps in a single subject. More realistic data analysis pipelines consist of many more steps in which often the same (or similar) pipelines are used for multiple subjects. In this section, we will show how the reproducescript functionality applies in such a case.

##### Original analysis

The analysis example follows the strategy outlined in (31) and starts with a single subject analysis pipeline that is repeated for four subjects. The directory is structured as depicted in figure 3. After the single-subject analysis, all single-subject results are used in a group analysis. The single-subject and group analyses are executed from the master script *analyze.m* (listing 4), which is the control script from which the relevant analysis scripts and functions are called. The master script relies on two functions: *doSingleSubjectAnalysis* and *doGroupAnalysis*, which are each stored in separate m-files. The original source code for these scripts can be found in Appendices Ia (single subject analysis) and IIa (group analysis).

**Figure 3.**
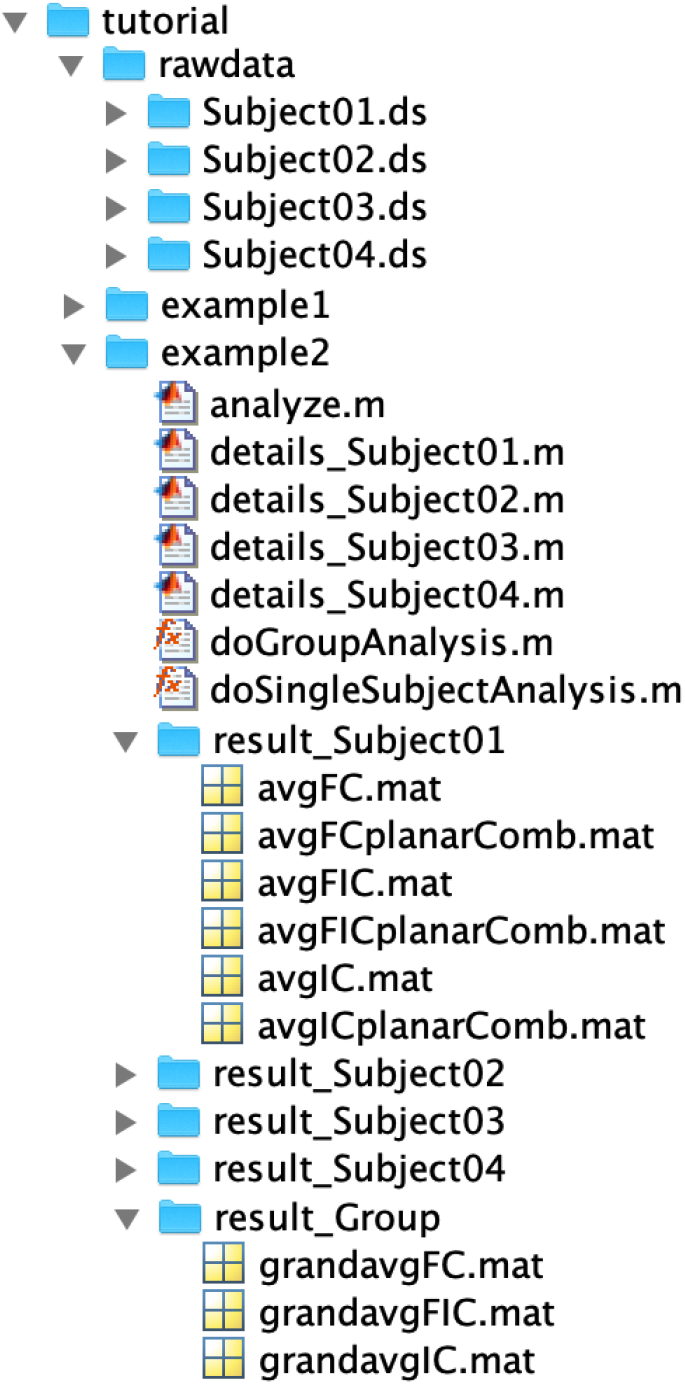
File directory tree for group study example. The main folder example2 contains scripts with source code and subject-specific analysis details, and separate folders for the results of each subject, and the group results for the original analysis (*result_\**).

**Listing 4.**
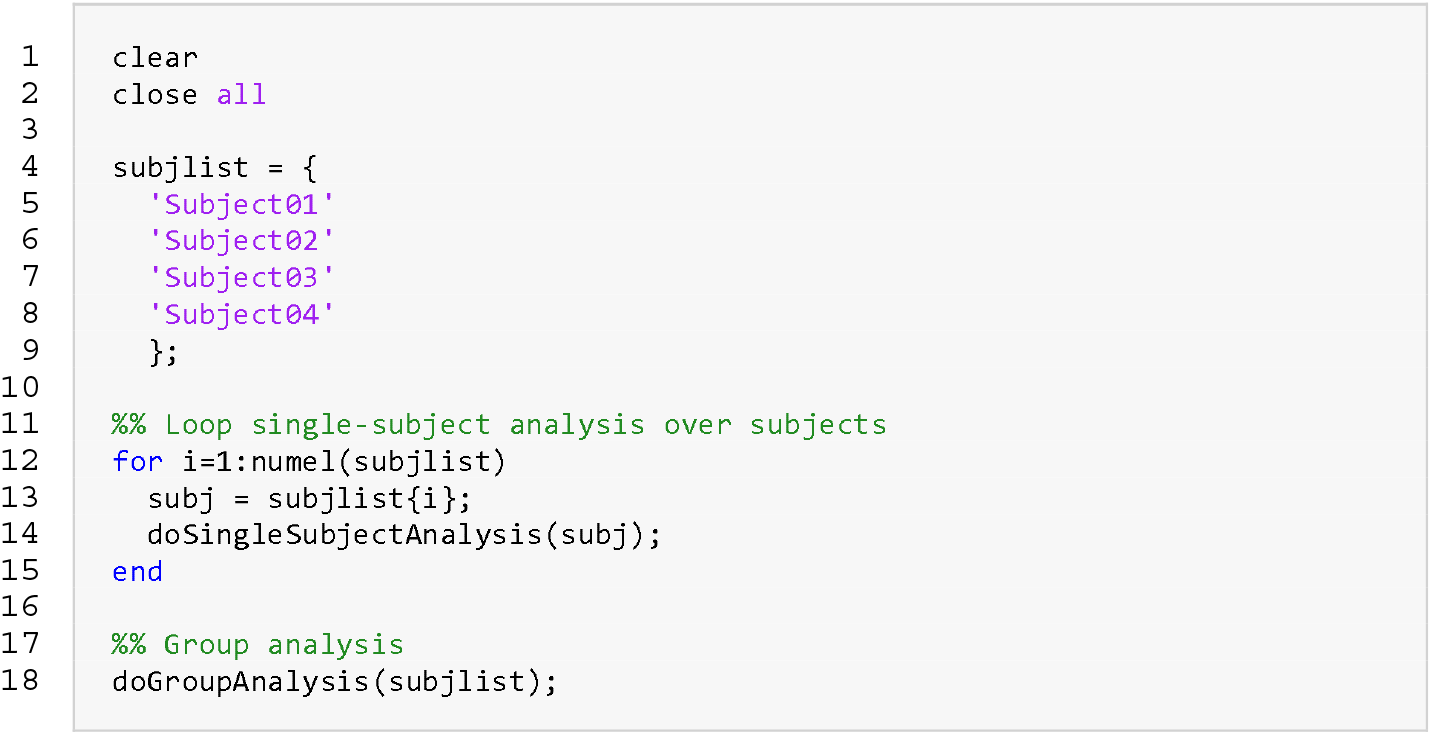
The entire group analysis can be executed from this master script.

### Initialization of reproducescript

To create a standard script from the analysis pipeline, the *ft_default* variable is initialized at the top of *analyze.m*. Note that we do not immediately initiate *reproducescript*, this is done in the loop just before *doSingleSubjectAnalysis*, and just before *doGroupAnalysis* by specifying unique directories (listing 5, lines 20 and 27) for each subject and foir the group. In fact, *reproducescript* can be stopped and restarted between different subjects, or even in between analysis steps, which is especially convenient in pipelines that require a lot of compute resources and that the researcher rather splits up to allow for parallel execution on a compute cluster.

**Listing 5.**
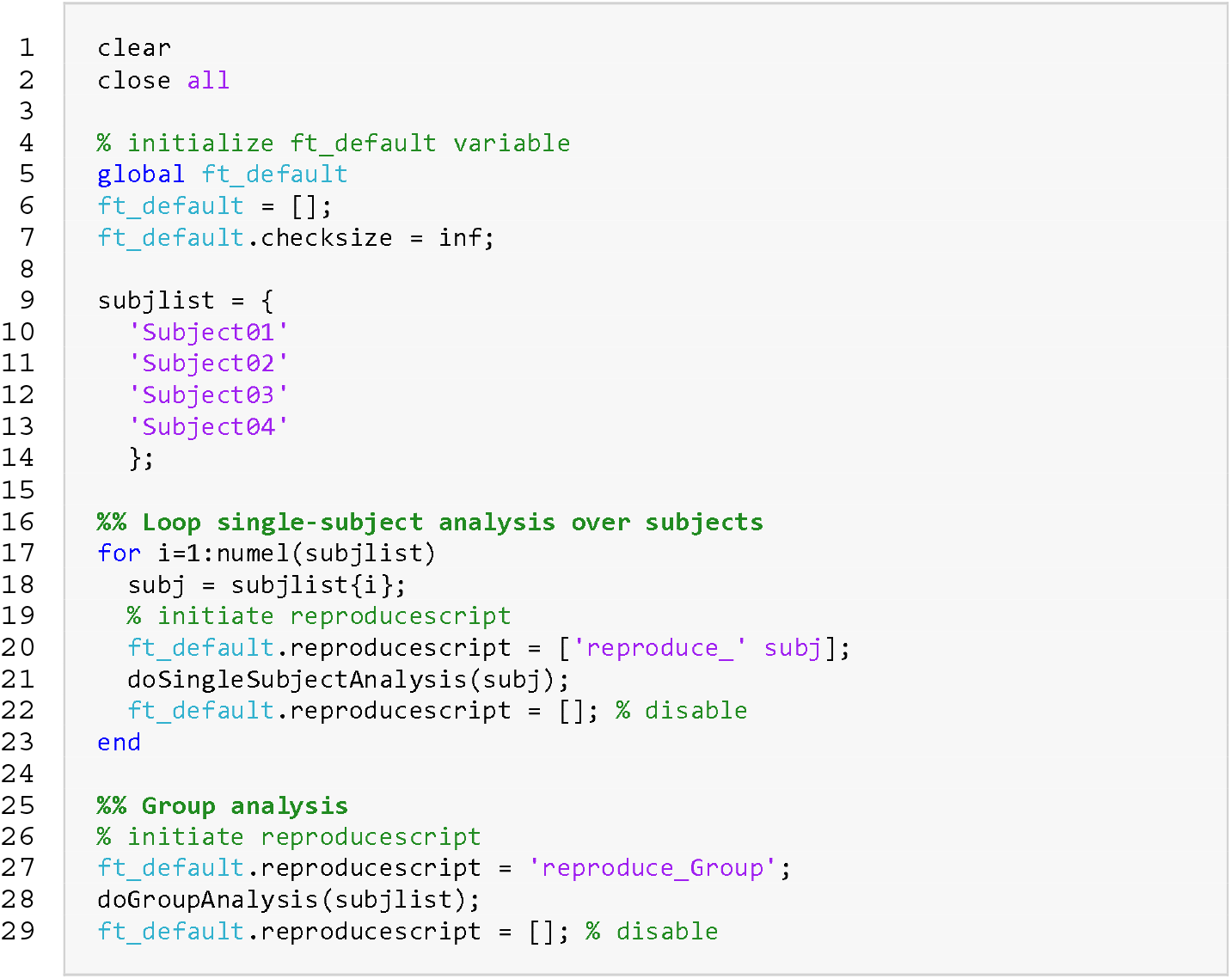
The master script from example 2 (listing 4), but now including the initialization of *reproducescript*.

### Reproduced analysis

The file directory tree (figure 4) and the initialization of *reproducescript* (listing 5) show that there is a specific folder devoted to the reproducescript content of each subject, and one for the group analysis. Thus, upon execution of the master script in listing 5, folders are created for each of the subjects, and for the group analysis. These all contain the intermediate data, a standardized script, and a *hashes.mat* file for the bookkeeping. The *reproducescript* standardized scripts for the single-subject analysis and group analysis can be found in the Supplementary Materials as Appendix Ib and IIb, respectively.

**Figure 4.**
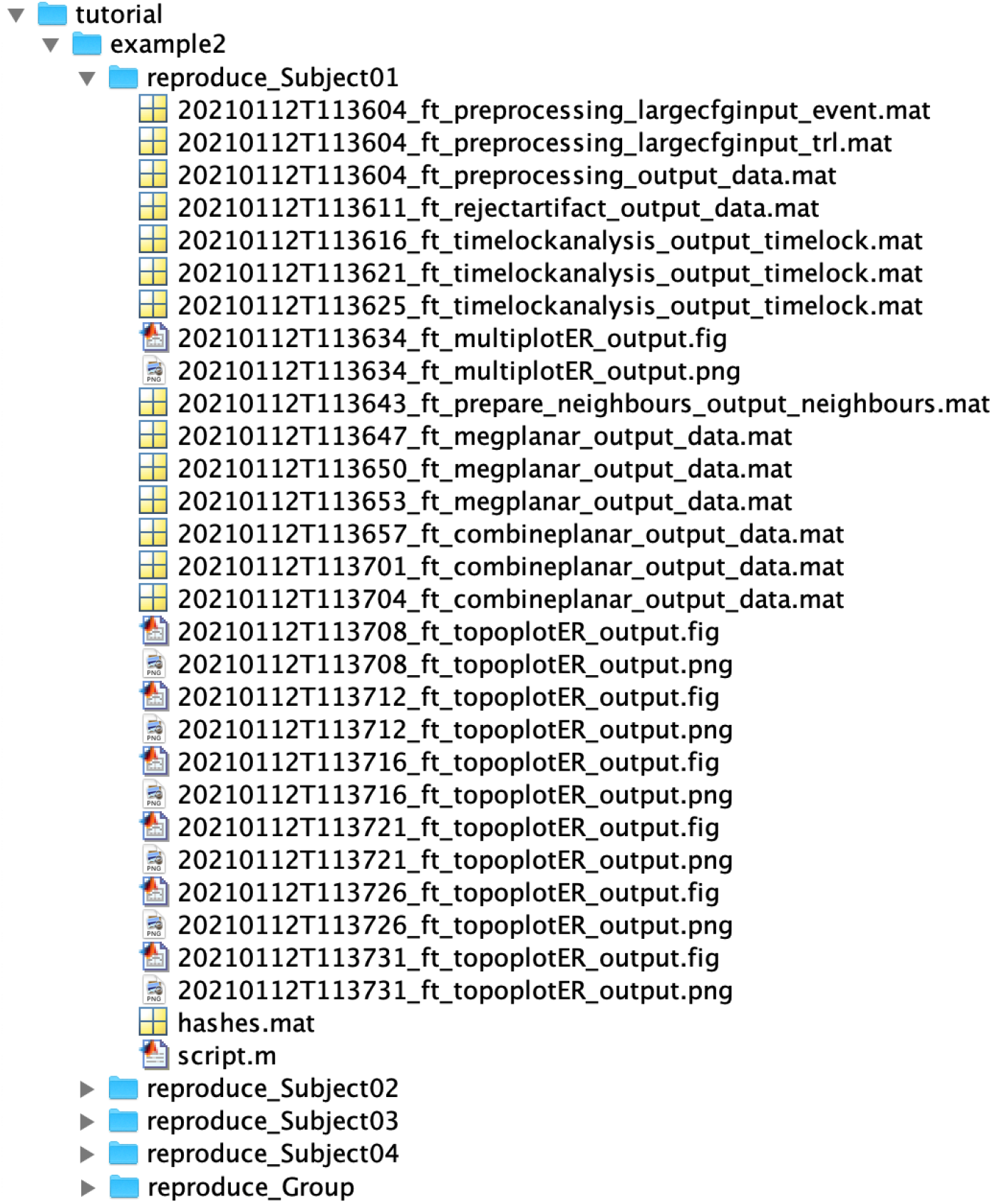
A *reproduce_\** folder is created for the reproduction of each single-subject analysis, and for the group analysis, containing the relevant (intermediate) data, standardized script, and hashes file for that analysis.

### Reproducescript in practice

In order to show that the reproducescript functionality is not limited to small examples and can indeed fully reproduce real-world pipelines with a single button click, we applied it to a previously published study that none of the authors of the current paper was involved in. The analysis pipeline in question was described by Andersen in (22) and published in a special Frontiers issue on group analyses on MEG data.

#### Example 3: application to published dataset

The analysis pipeline in (24) is well-documented and itself a good demonstration of a reproducible analysis pipeline in the FieldTrip ecosystem. Nevertheless, it consists of a complex set of 10 analysis scripts and 46 functions, which, without the extensive documentation that has been provided by the author, would be challenging to reuse and reproduce the results. This makes it particularly suited to demonstrate the effectiveness and simplicity of *reproducescript*.

Andersen describes an analysis pipeline from raw single-subject MEG data to group-level statistics in source space. Each of the custom-written scripts has a specific purpose (figure 5), but multiple analysis steps in separate functions are required for the purpose of one script (see figure 6 for the full analysis pipeline), creating a complex hierarchy of scripts and functions.

**Figure 5.**
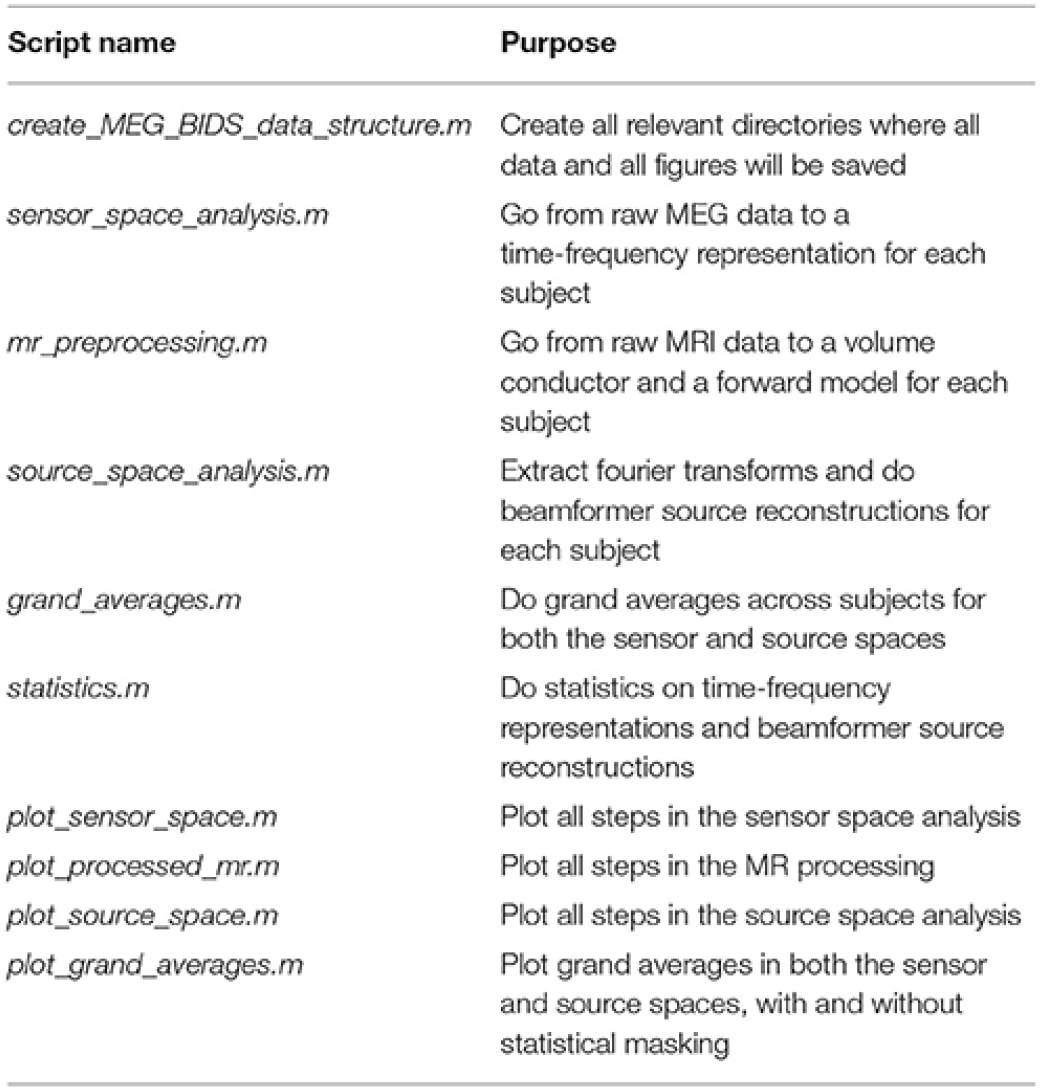
The purpose of each original script, reproduced with permission from (24).

**Figure 6.**
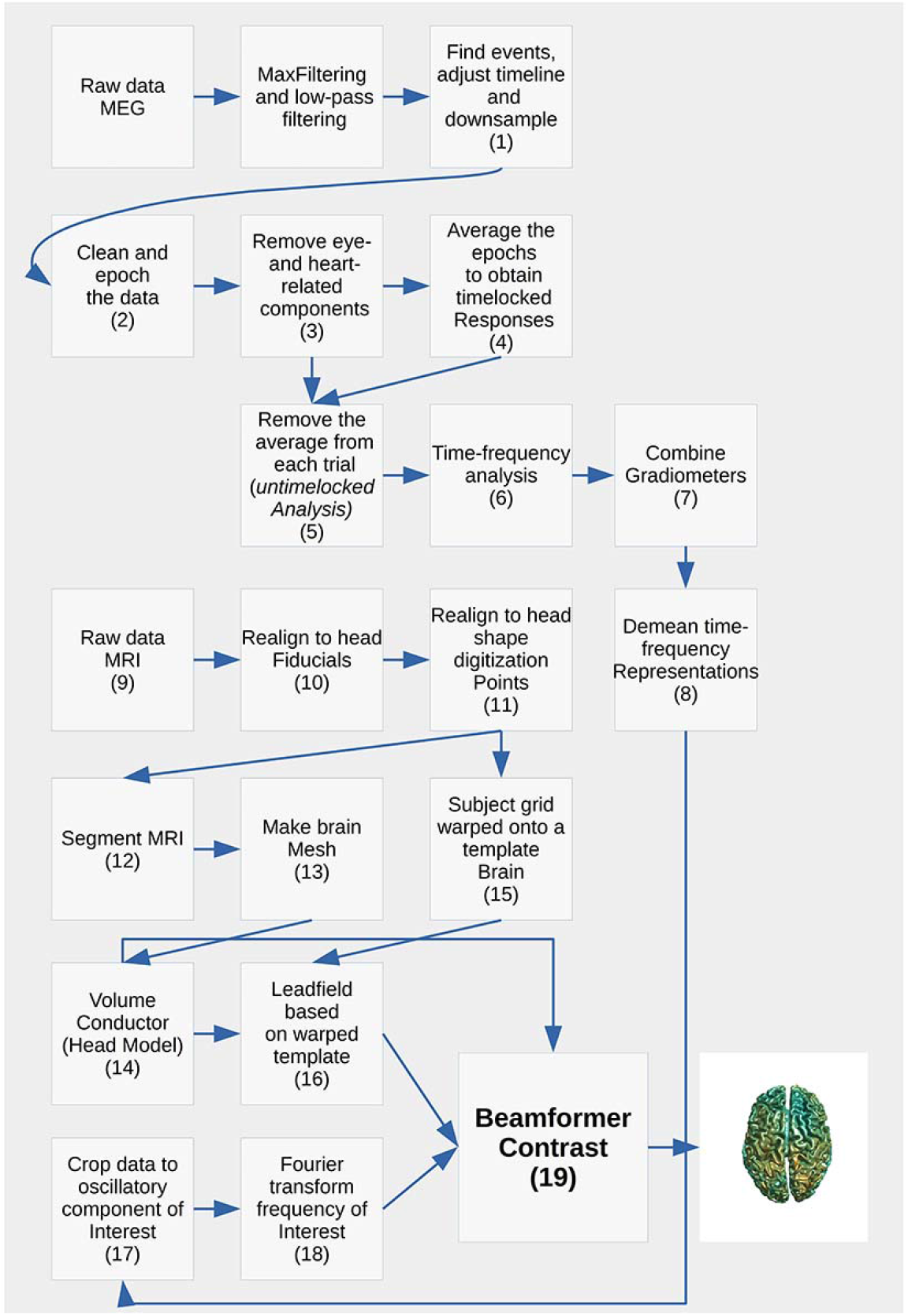
The full analysis pipeline (reproduced with permission from (24)) does not linearly map onto a single custom-written script or function.

To keep the computational time and storage requirements low, we applied the full analysis pipeline to two subjects only. The pipeline was ran with *reproducescript* enabled, and thus created the original results, and also the code and intermediate data with which it should be able to reproduce the results, had the original scripts not been available. Both the original source code from Andersen and the standardized scripts generated by *reproducescript* are available on GitHub (https://github.com/matsvanes/reproducescript). To confirm that *reproducescript* indeed resulted in a reproducible analysis pipeline, the newly formed standardized script was executed, and its results were compared qualitatively (figures) and quantitatively (data) with the original results.

Even though the original source code from Andersen is organized clearly and is accompanied by extensive documentation, it did take some effort to get the analysis pipeline up and running. Besides changing the relevant directories in the analysis scripts, the initial executions of the pipeline resulted in unexpected errors (e.g. some data files were missing because they contained information that could trace back to a specific individual; without these files, the pipeline broke). This illustrates that even the cleanest hand-written analysis pipelines might not be easily reproducible. After all errors were resolved (i.e. through help of the corresponding author of the original pipeline) the pipeline was executed with *reproducescript* enabled, similarly to the group study in the example above (see the superscript in listing 6).

The resulting data that was produced by the original pipeline applied to the two subjects amounted to roughly 25 GB of data (± 12 GB per subject and 1 GB for group analysis) and a similar amount in the figures. The reason that the figures make up a lot of data is that they are saved both as .png and as .fig files. The .png files are small because they only contain pixel values of the image, while .fig files contain the complete data that was plotted (in compressed format), and thus these files are large.

Because *reproducescript* saves all intermediate data, the total amount of data was higher: 140 GB (± 64 GB per subject and 12 GB for group analysis), and 18 GB of figures. If we extrapolate these numbers to the entire group study (e.g. 20 subjects) and save figures as png, the original pipeline would result in ± 240 GB of data, and the reproducescript version in roughly 1300 GB, or 5.4 times the disk space requirements of the original pipeline. Note that this is an example and by no means a rule of thumb. The amount of data produced by reproducescript will vary between pipelines and depends on the amount of FieldTrip calls. The *reproducescript* pipeline will amount to more data than the original in almost all cases, because all intermediate data is saved, which is typically not done in original analysis scripts.

The *reproducescript* analysis pipeline was executed without further problems and without the need for debugging. We asserted whether this pipeline produced the same results as the original pipeline. In the group analysis, the last analysis step comprises a statistical comparison between two conditions, using *ft_frequencystatistics* and *ft_sourcestatistics*. The results from both pipelines were numerically identical. Even if a FieldTrip function relies on random numbers (e.g. for the initialization of an ICA algorithm, or for a random permutation in statistics), numerically identical numbers can be acquired: if a FieldTrip function uses random numbers, *reproducescript* saves the state of the random number generator in the standardized script. This allows the exact numerical results to be obtained from the *reproducescript* pipeline. However, this is not always to be expected, especially when interactive and subjective analysis steps are part of the analysis. For example, during preprocessing a researcher could visually select particular trials and/or channels with artifacts and reject them from the data. However, a different researcher might employ other criteria and thus remove other pieces of data as artifacts. Therefore, it is not always expected (or desired) to get numerically identical results from the reproduced pipeline (see Discussion).

The comparison between the original pipeline and the reproduced pipeline revealed a few more caveats. First, only analysis steps that were conducted in the FieldTrip ecosystem can be reproduced, since source code outside of the FieldTrip ecosystem is not tracked by *reproducescript*. In the current example, Andersen used several standard MATLAB plotting functions for visualization of the results, which were not reproduced by the *reproducescript* pipeline. This can especially be problematic when these steps are the last in the pipeline and represent the outcome of the analysis. There are options to work around this issue. For example, the user can provide extra documentation in the standardized script, describing how the figures can be reproduced manually. The same holds for transformations of the data outside the FieldTrip ecosystem. Instead of writing comments in the standardized script after running the analysis pipeline, the researcher could also use *ft_annotate* in the original code after using a function outside the FieldTrip ecosystem. This returns the same output data as the user has provided as input, but allows the researcher to add comments to that data structure, which then become part of the provenance that is stored in the data structure. Another option is to embed non-FieldTrip code into a copy of *ft_examplefunction*, which contains all essential (‘boilerplate’) FieldTrip bookkeeping functionality, and thereby wrap the original custom code in a new FieldTrip function. Both these options require some extra time investment of the researcher. Therefore, the more the pipeline relies on functions within the FieldTrip ecosystem, the less work to make the pipeline reproducible and transparent.

Second, even if the pipeline exclusively uses FieldTrip functions, some FieldTrip functions evaluate custom-written code. For example, a user can specify custom code to select trials in *ft_definetrial* (i.e. *cfg.trialfun*). If this code were not shared, this particular analysis step could not be re-executed, but since intermediate results are stored as well (in the example of *cfg.trialfun*, *cfg.trl* is stored), it is always possible to skip a particular step and continue with the rest of the pipeline.

**Listing 6.**
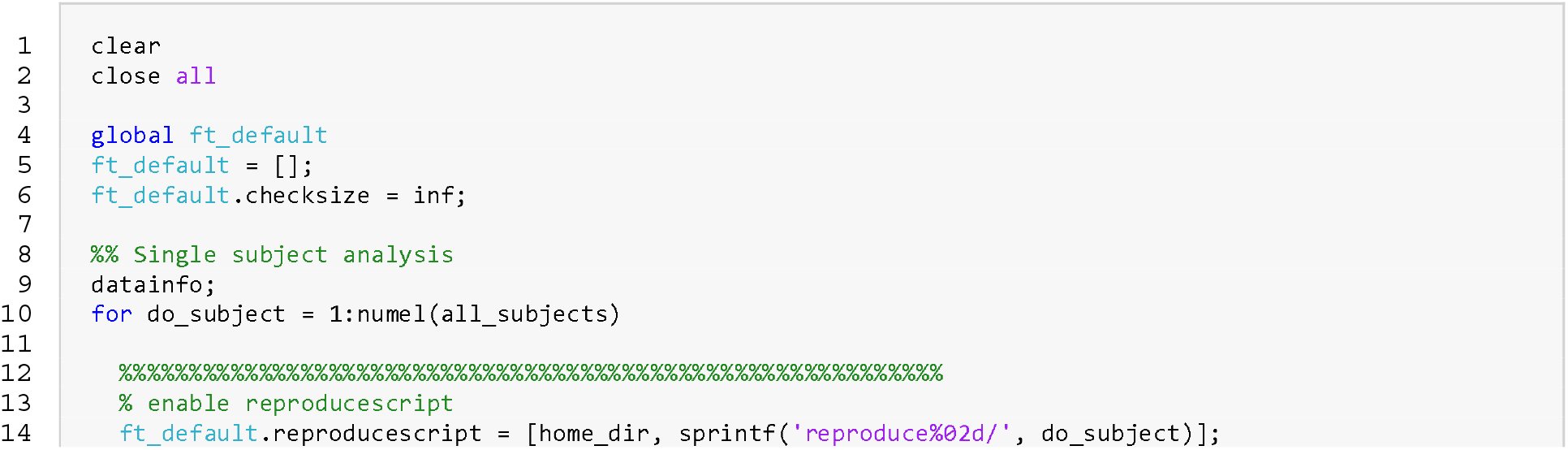
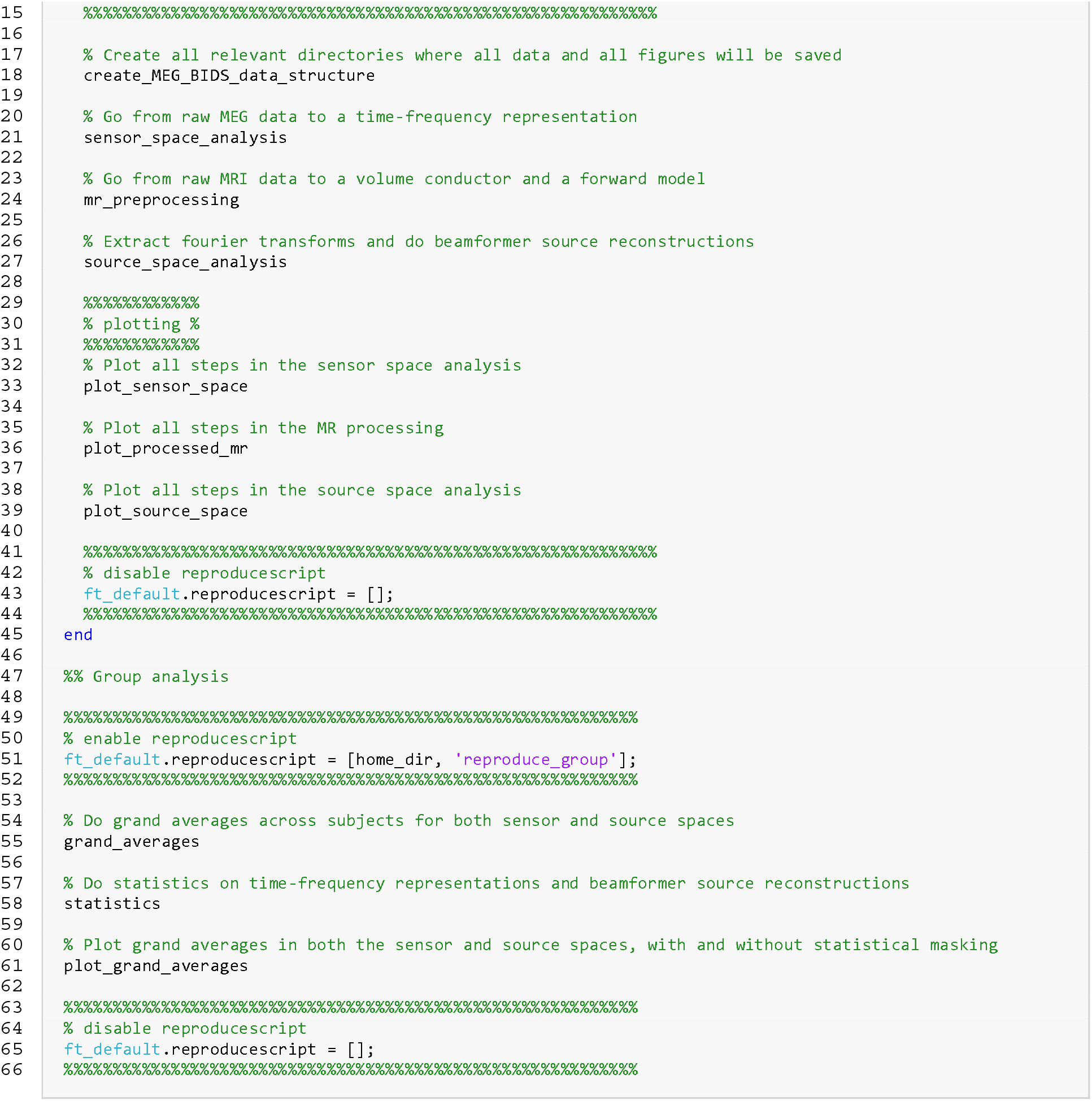
Master script for running the pipeline of Andersen (24) with *reproducescript* enabled. This master script was not explicitly part of the source code shared by Andersen, but was created based on his documentation.

## Discussion

Neuroimaging research is relying more and more on complex computational analysis pipelines. Furthermore, there are strong motivations to improve the reproducibility of neuroimaging results. Published results can in part be verified by having access to the details and being able to reproduce the analysis pipeline that produced them.

We presented the design and implementation of new functionality that can help individual researchers to generate an analysis pipeline that is generic, reproducible, and can be shared easily, while requiring only minimal effort on the researcher’s part. The functionality, termed *reproducescript*, has been released (May 2020) as part of the FieldTrip toolbox. It generates a standardized executable script from the original researcher’s source code, which can be complex, complicated, and/or of variable quality. By providing this tool, we hope to encourage researchers to share their analysis pipelines and corresponding data more commonly.

We verified the effectiveness of *reproducescript* by applying it to a published analysis pipeline (24). The example we applied it to is already a good example of a reproducible analysis pipeline, but contains a hierarchy of custom written scripts and functions on top of the FieldTrip toolbox. We re-executed the original published pipeline with *reproducescript* enabled, and found that the generated, standard-format, code was able to reproduce the original results faithfully.

### *reproducescript* reproduces analysis pipelines efficiently and transparently

The reproducible analysis pipeline was easy to execute and did not require debugging. The results of the reproduced pipeline were numerically identical to those of the original pipeline, which is a demonstration of the robustness of *reproducescript*. This is not to say that numerical identity should always be the goal of reproduction efforts. Instead of asking “did this code with these exact parameters return these exact numerical results”, it sometimes is more insightful to show that an analysis pipeline will return the same *qualitative* results, independent of arbitrary choices in preprocessing and the state of random number generators. With *reproducescript*, both are possible. If numerical reproduction is required, choices in interactive analysis steps can be based on the researcher’s documentation if provided, obtained from the input- and output- files of the interactive step. Otherwise, the interactive step can simply be skipped (i.e. the input- and output- data that are generated by the first run of the original pipeline are trivially identical to the original results). If only qualitative reproduction is desired, the state of the random number generator can be removed from the reproduced script (i.e. *cfg.randomseed*) and the researcher can run the entire pipeline.

In addition to providing quantitative or qualitative reproducibility for an analysis pipeline, *reproducescript* ensures that all analysis steps remain transparent: every individual analysis step is interpretable, and even though the standardized script might become large, the complete pipeline can easily be explored using a standard text editor or the visualization tools on GitHub, or visualized with *ft_analysispipeline*.

### Limitations of *reproducescript*

One drawback is that the presented method is limited to analysis steps that rely on functions within the FieldTrip ecosystem. Manual source code, or code that uses default MATLAB functions, are not tracked by *reproducescript* and those steps will therefore not be represented in the standardized script. This means that the transparency of those particular analysis steps is limited, and it is up to the researcher to provide documentation about what transformations of the data are applied in those steps. If a large part of the analysis pipeline does not use FieldTrip functionality, the benefits of *reproducescript* will be limited. Therefore, this functionality is mostly targeted at researchers who do not use a lot of custom written code or external toolboxes, with the important exception that custom control structures and functions wrapping FieldTrip functionality are tracked. As in the real-life example we presented here, *reproducescript* will faithfully track any custom functions, loops, etc. written by the researcher that internally make use of FieldTrip functionality; and, in fact, we would recommend researchers to use such control structures when building their analysis pipelines. Researchers who tend to use large amounts of custom written code to do actual analysis work - as opposed to merely structuring the flow of the code - are probably also those with more expertise in writing and documenting source code, and are therefore already better equipped to produce a reproducible analysis pipeline or adapt their code such that it becomes part of FieldTrip’s provenance (e.g. by building it into *ft_examplefunction*).

## Conclusion

In conclusion, we believe to have provided researchers with a tool to easily share complete analysis pipelines that use the FieldTrip toolbox.

This tool is not meant to replace already existing solutions, like pipeline systems, version control systems and literate programming, which we think have great value. However, in certain cases, these tools are limited in their functionality (e.g. the rigidity of pipeline systems) or they require a lot of time investment (learning to use version control systems or writing analysis code in a literate programming style). While *reproducescript* is limited in its use for those researchers whose analysis pipelines rely largely on custom written algorithms or external toolboxes, it can be of great use for the large group of researchers who mostly or even exclusively use FieldTrip. *reproducescript* can be used flexibly, and it can reproduce results both quantitatively and qualitatively, all the while keeping the pipeline transparent and intuitive. All of this can be done without much effort by either the researcher providing the pipeline or the researcher executing it. By making it easier for researchers to share their reproducible analysis pipeline, we hope this functionality will help to make science more robust and transparent.

## Data and code Availability

*reproducescript* is part of the open source FieldTrip toolbox, which can be downloaded from https://www.fieldtriptoolbox.org. The raw data for example 1 and 2, and the output data from all examples can be accessed via the Donders Repository (https://doi.org/10.34973/21pa-dg13). The raw data from example 3 was described by Andersen (24) and can be accessed at https://doi.org/10.5281/zenodo.1134776. All MATLAB scripts for the analysis of the data and reproduction of the results are available at https://github.com/matsvanes/reproducescript.

## Author Contributions

Mats W.J. van Es

Roles: Conceptualization, Data Curation, Formal Analysis, Investigation, Methodology, Project Administration, Resources, Software, Validation, Visualization, Writing – Original Draft Preparation, Writing – Review & Editing

Eelke Spaak

Roles: Conceptualization, Data Curation, Funding Acquisition, Methodology, Resources, Software, Writing – Review & Editing

Jan-Mathijs Schoffelen

Roles: Conceptualization, Data Curation, Funding Acquisition, Methodology, Resources, Software, Writing – Review & Editing

Robert Oostenveld

Roles: Conceptualization, Data Curation, Funding Acquisition, Methodology, Resources, Software, Supervision, Writing – Review & Editing

## Competing interests

There are no competing interests to be disclosed

## Funding Information

This work was supported by The Netherlands Organisation for Scientific Research, NWO Vidi: 864.14.011 awarded to Jan-Mathijs Schoffelen, NWO Veni: 016.Veni.198.065 awarded to Eelke Spaak. The funding source had no role in the realization of this manuscript.

## Acknowledgements

The authors would like to thank Lau Andersen for publishing his original data and analysis scripts in (24) and his help in executing the pipeline.

## Supplementary Material

## Appendix Ia Group study – single subject analysis

Appendix Ia lists the (hand-written) code used in the single-subject analysis of example 2 in the main text.

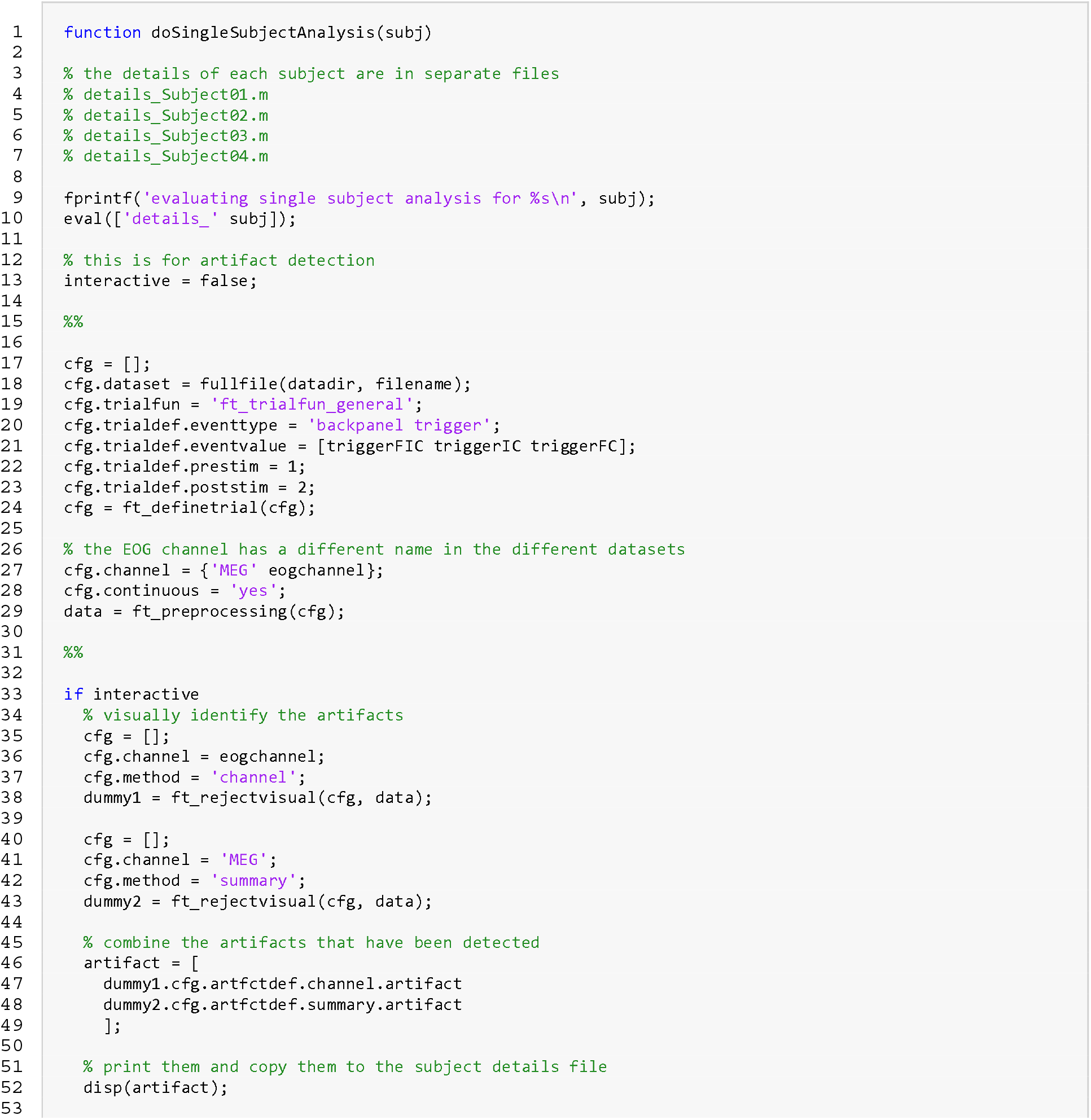
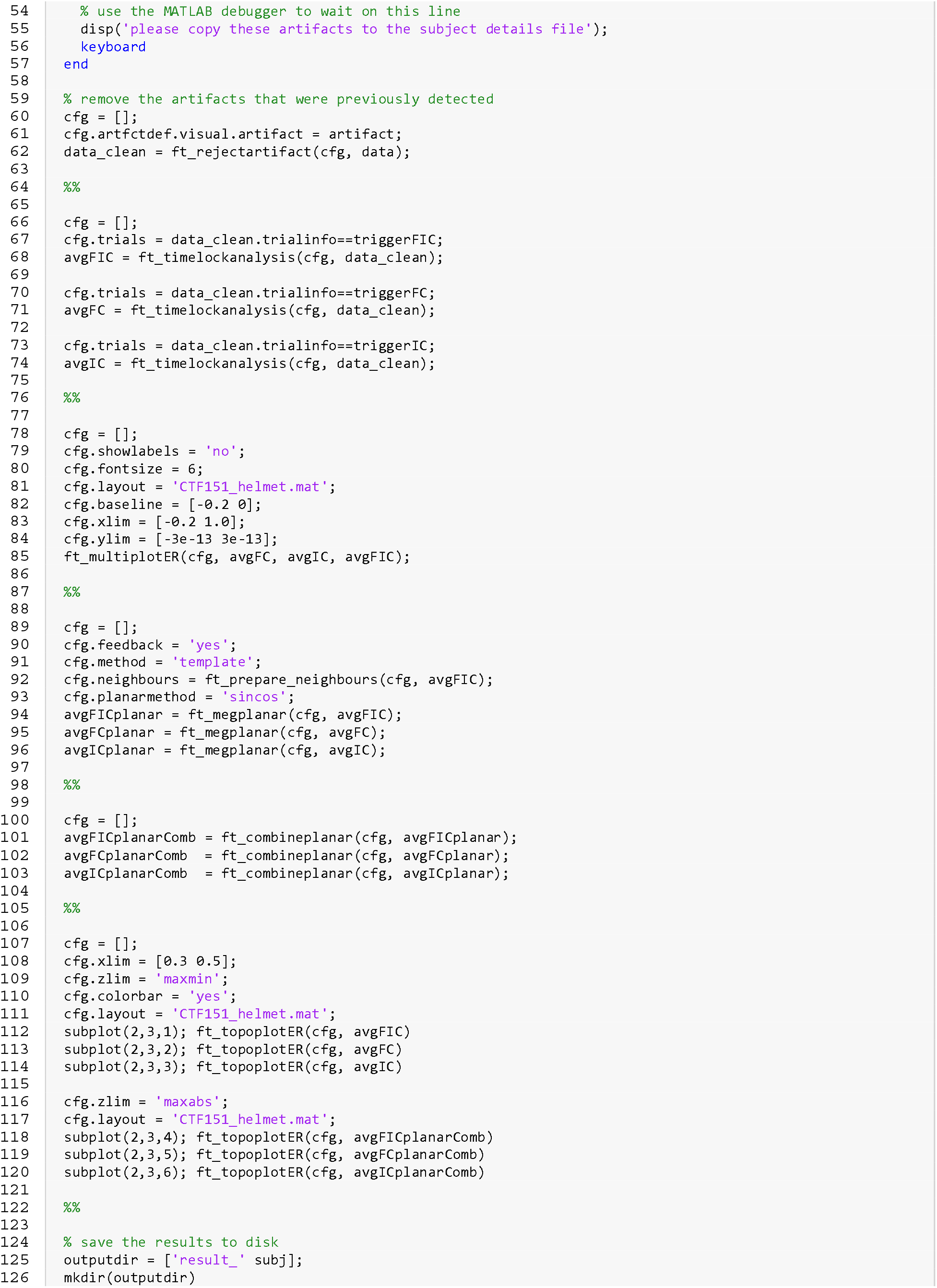
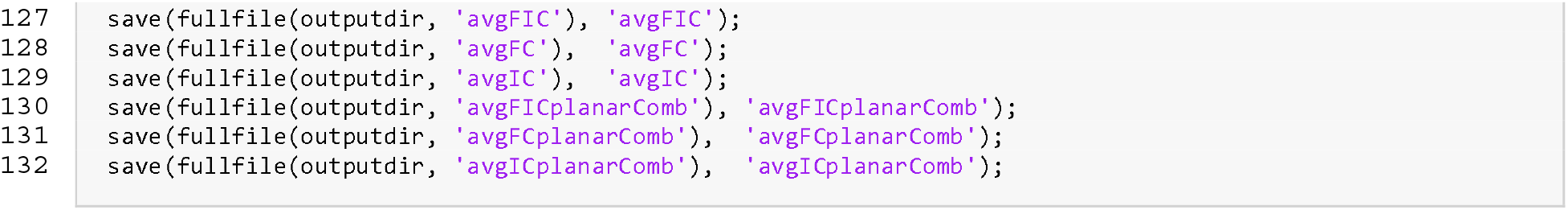

## Appendix Ib Group study – single subject analysis (reproducescript)

Appendix Ib lists the code for the single-subject analysis produced by reproducescript after running example 2 from the main text. It is the *reproducescript* counterpart of the source code in Appendix Ia, executed for Subject01.

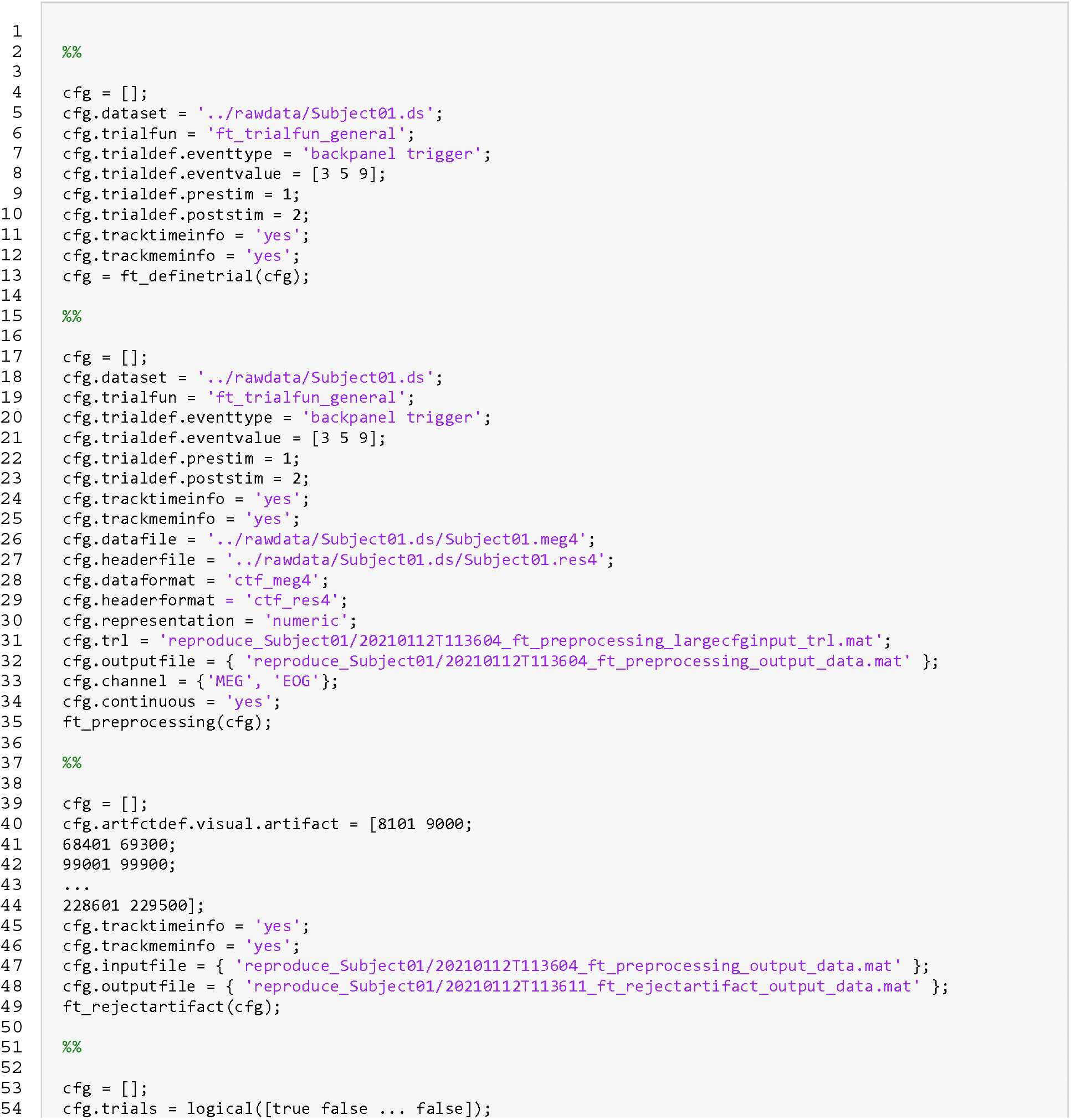
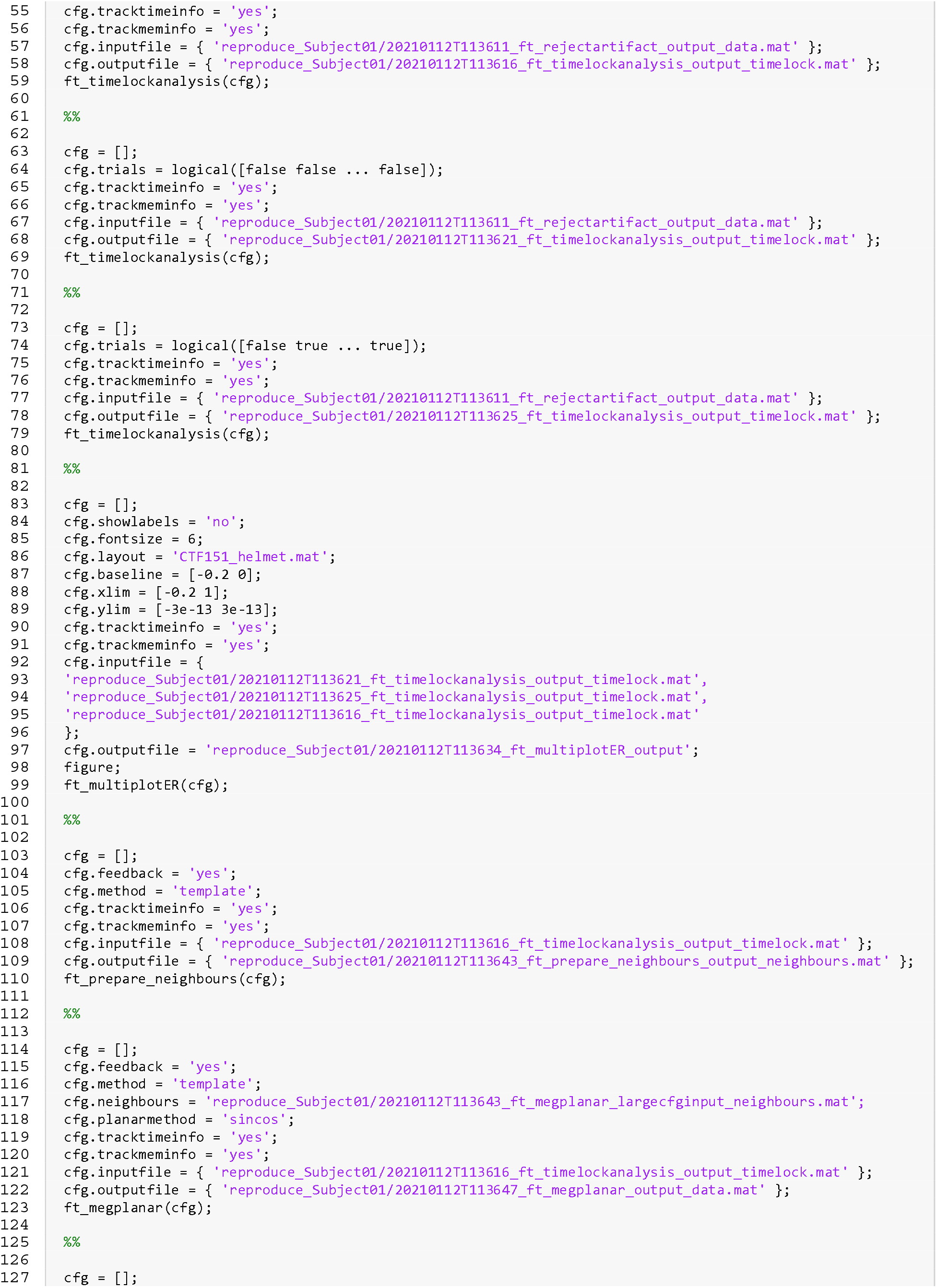
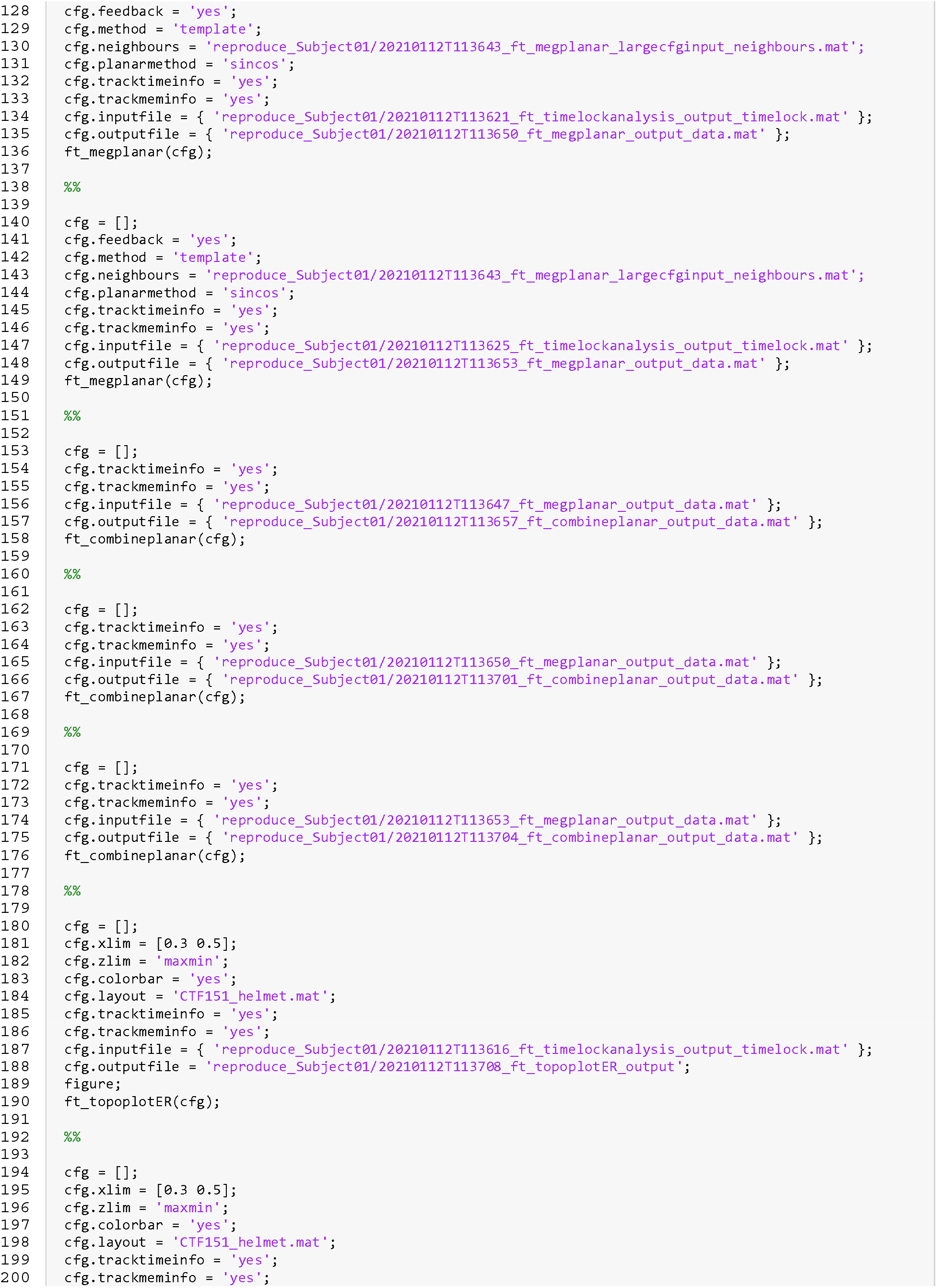
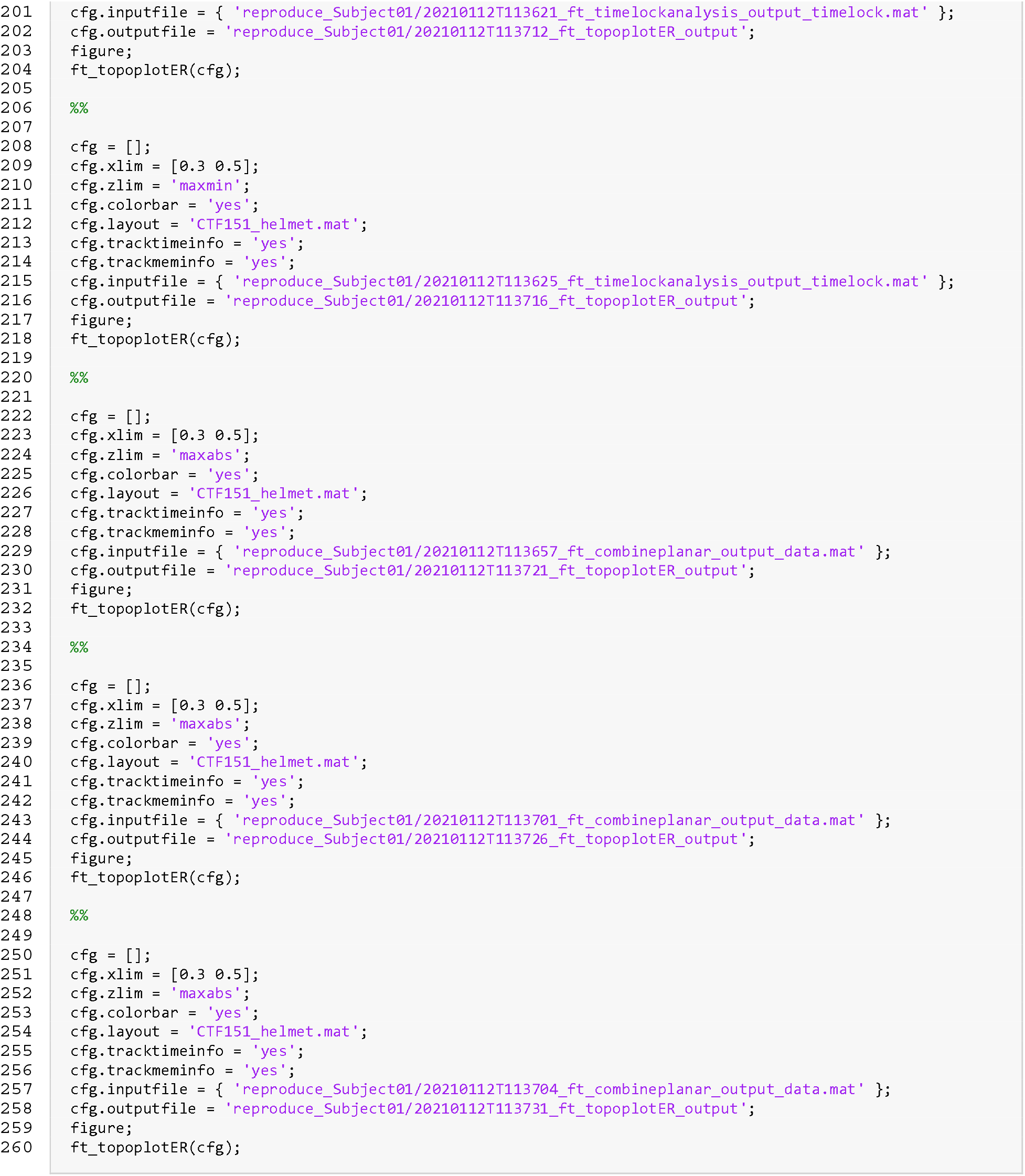

## Appendix IIa Group study – group analysis

Appendix IIa lists the (hand-written) code used in the group analysis of example 2 in the main text.

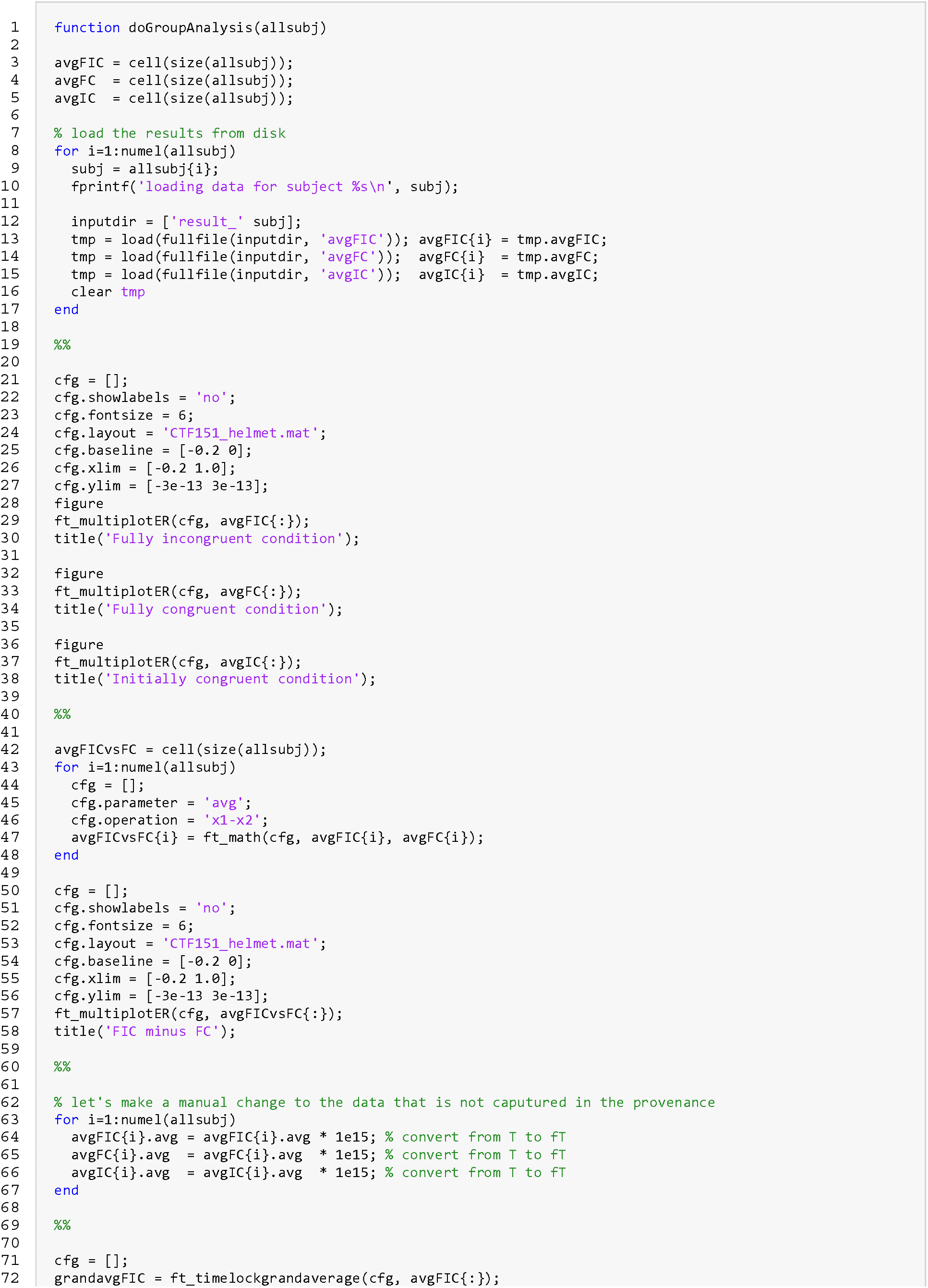
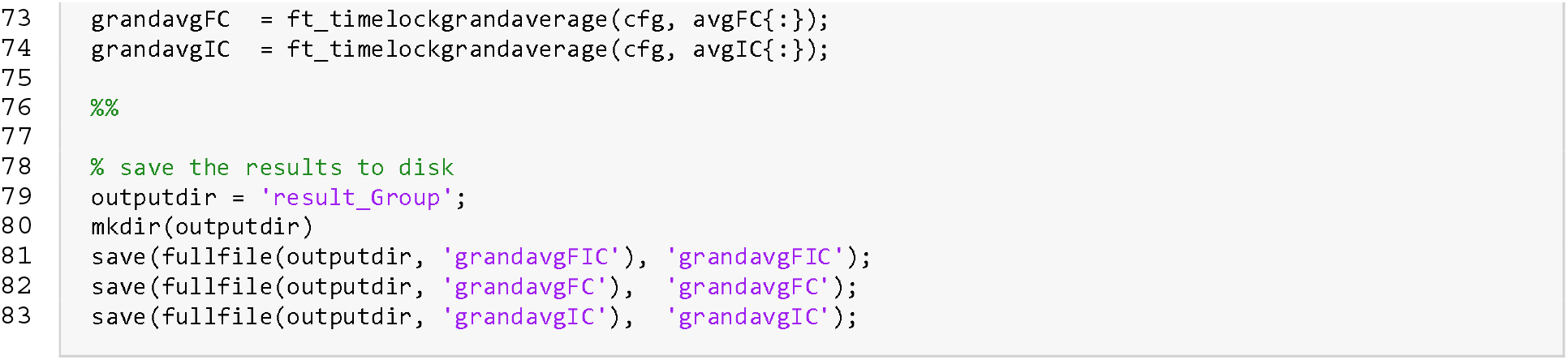

## Appendix IIb Group study – group analysis (reproducescript)

Appendix IIb lists the code for the group analysis produced by *reproducescript* after running example 2 from the main text. It is the *reproducescript* counterpart of the source code Appendix IIa.

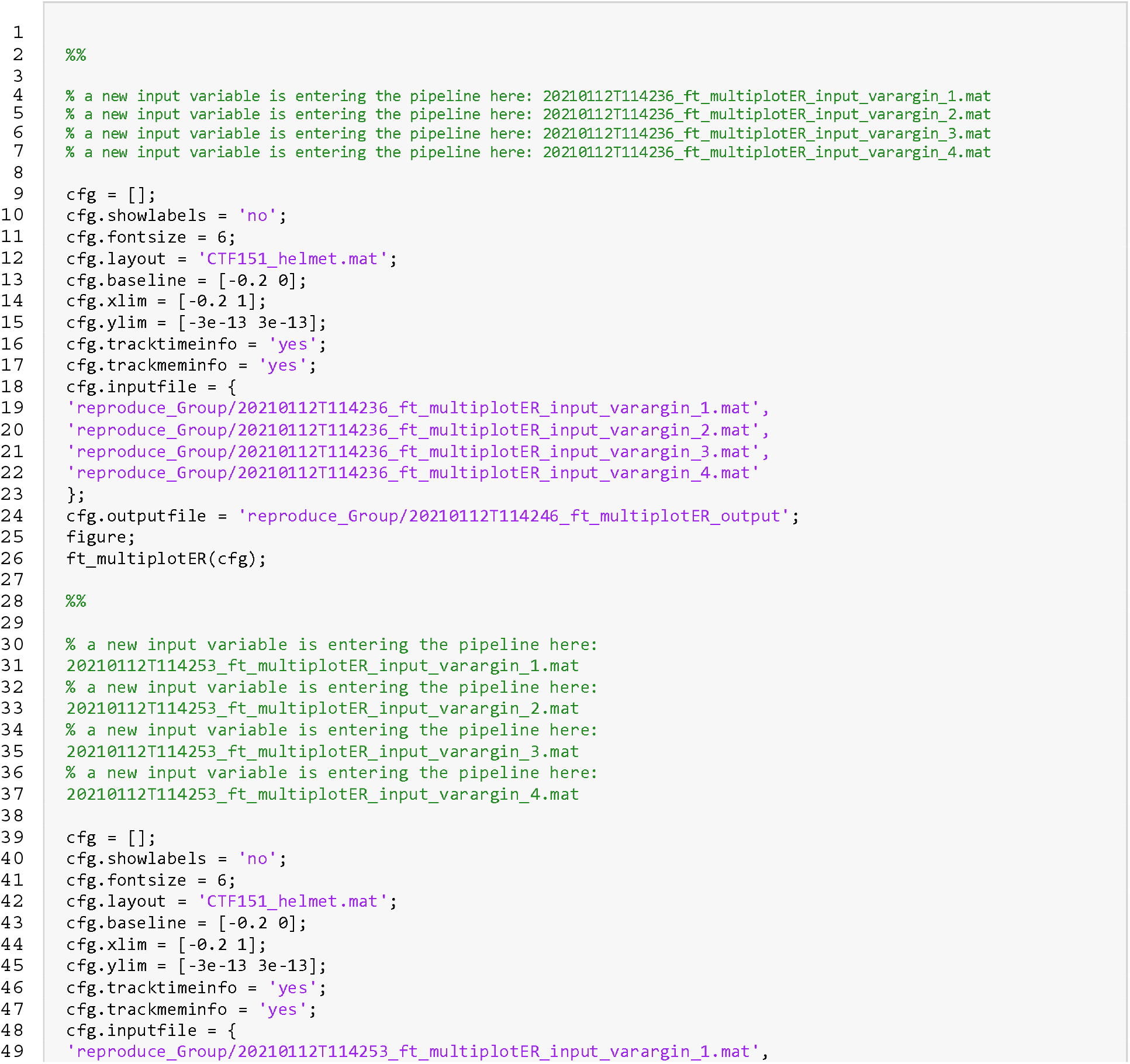
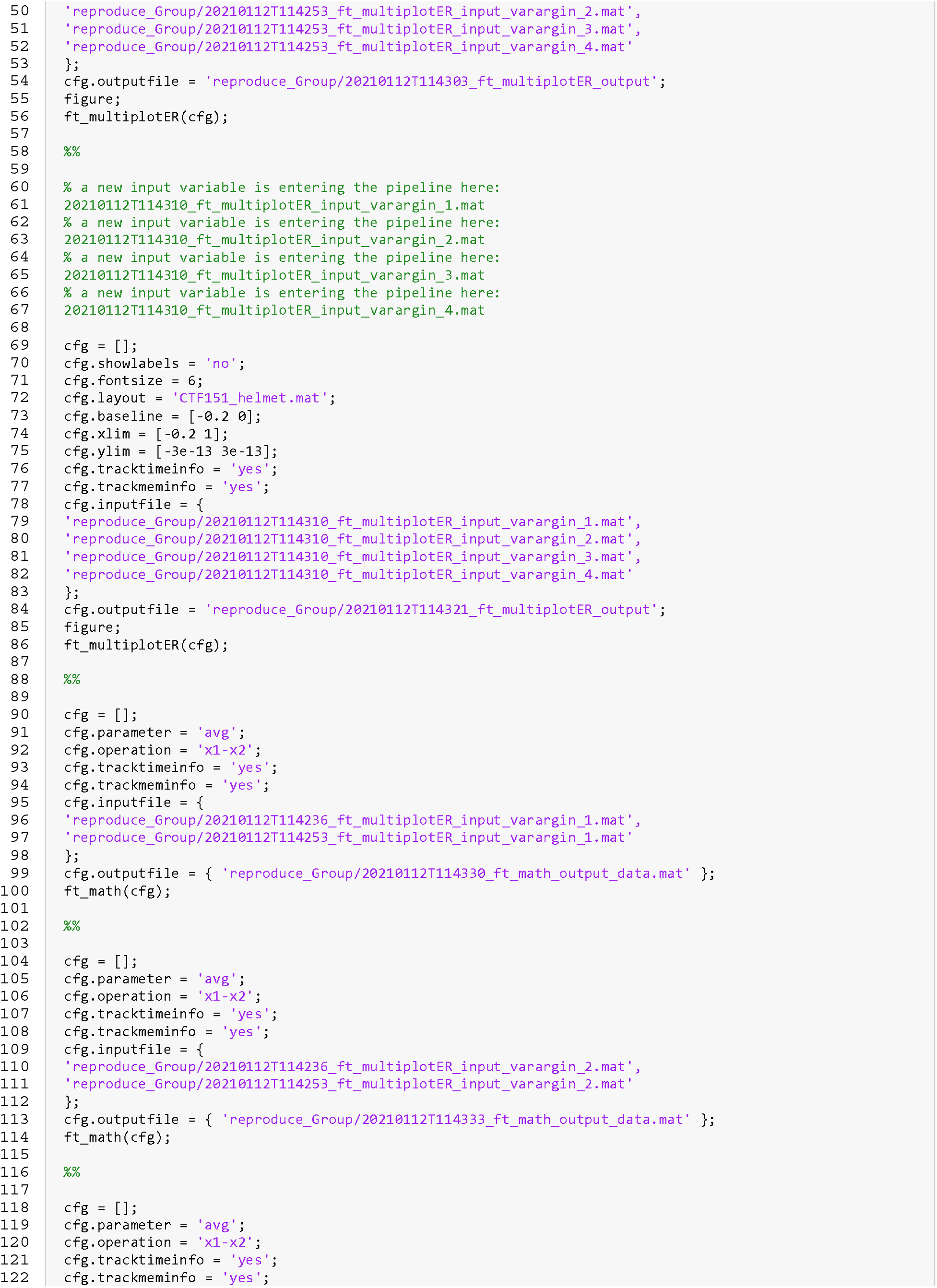
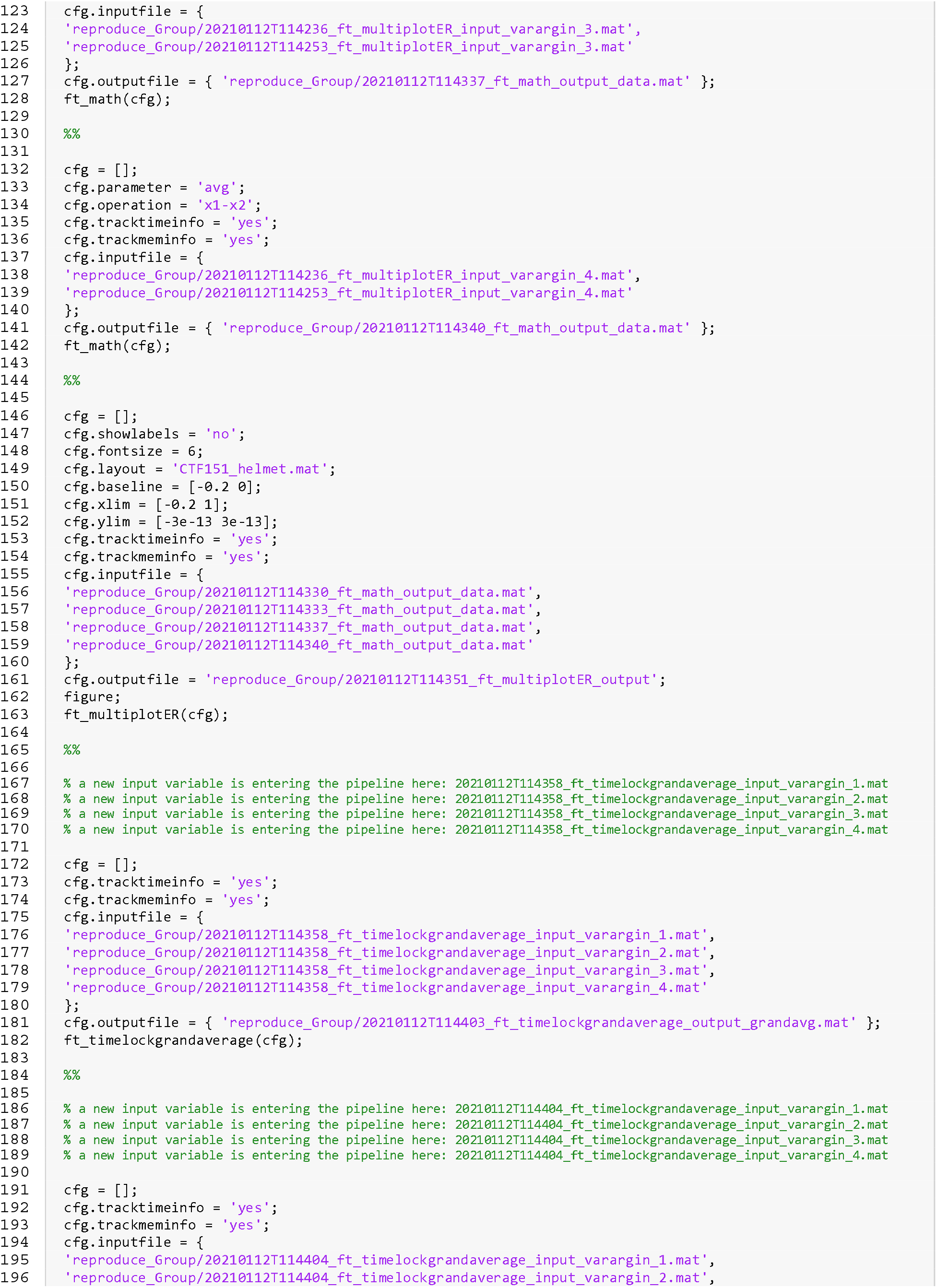
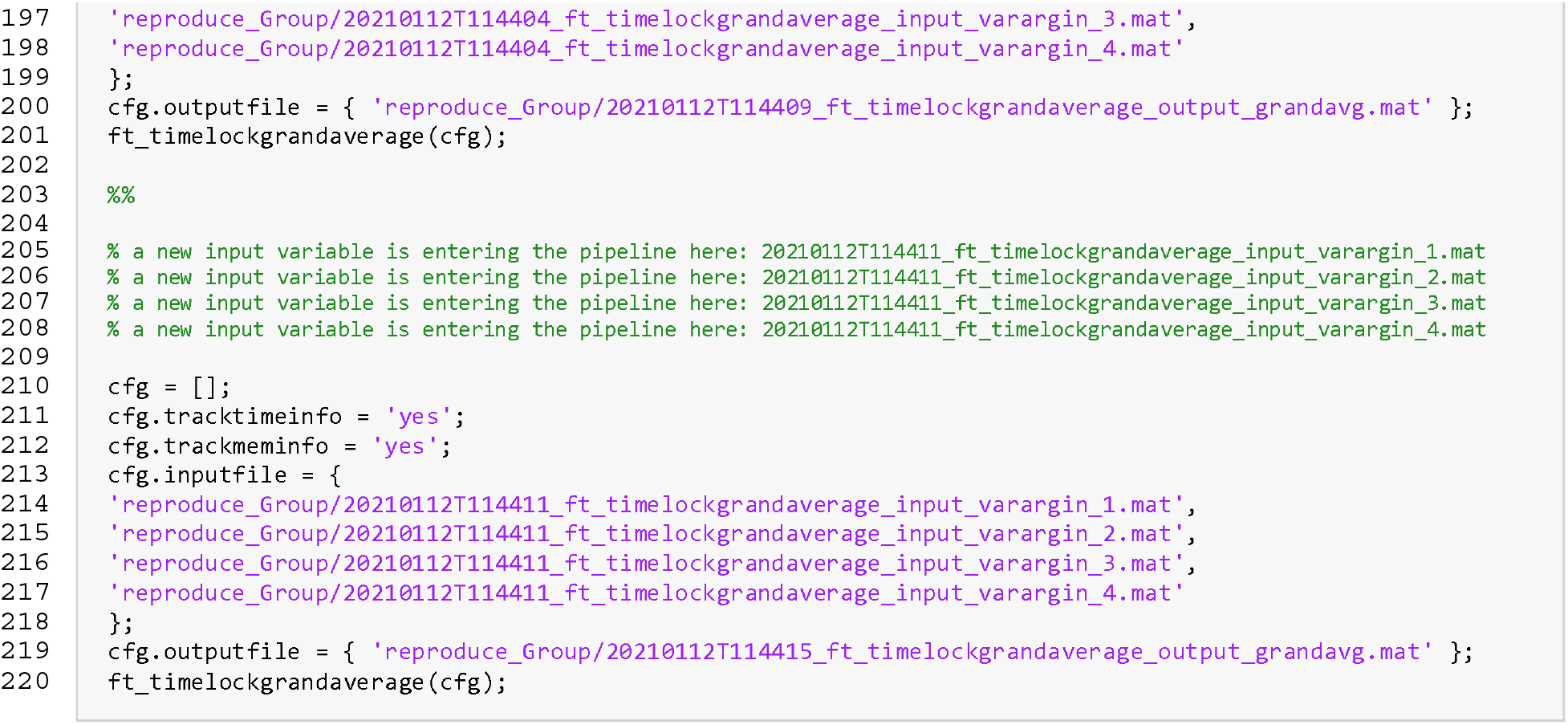

## Notes

https://www.fieldtriptoolbox.org/

https://doi.org/10.34973/21pa-dg13

https://doi.org/10.5281/zenodo.1134776

https://github.com/matsvanes/reproducescript

### Competing Interest Statement

The authors have declared no competing interest.

## References

1. Open Science Collaboration. Estimating the reproducibility of psychological science. Science. 2015 Aug 28;349(6251):aac4716–aac4716.

2. Simmons JP, Nelson LD, Simonsohn U. False-Positive Psychology: Undisclosed Flexibility in Data Collection and Analysis Allows Presenting Anything as Significant. Psychol Sci. 2011 Nov;22(11):1359–66.

3. Button KS, Ioannidis JPA, Mokrysz C, Nosek BA, Flint J, Robinson ESJ, et al. Power failure: why small sample size undermines the reliability of neuroscience. Nat Rev Neurosci. 2013 May;14(5):365–76.

4. Gilmore RO, Diaz MT, Wyble BA, Yarkoni T. Progress Toward Openness, Transparency, and Reproducibility in Cognitive Neuroscience. Ann N Y Acad Sci. 2017 May;1396(1):5–18.

5. Szucs D, Ioannidis JPA. Empirical assessment of published effect sizes and power in the recent cognitive neuroscience and psychology literature. PLoS Biol [Internet]. 2017 Mar 2 [cited 2020 May 6];15(3). Available from: https://www.ncbi.nlm.nih.gov/pmc/articles/PMC5333800/

6. Gleeson P, Davison AP, Silver RA, Ascoli GA. A Commitment to Open Source in Neuroscience. Neuron. 2017 Dec;96(5):964–5.

7. Zwaan RA, Etz A, Lucas RE, Donnellan MB. Making Replication Mainstream. Behavioral and Brain Sciences. 2017 Oct 25;1–50.

8. Gross J, Baillet S, Barnes GR, Henson RN, Hillebrand A, Jensen O, et al. Good practice for conducting and reporting MEG research. NeuroImage. 2013;65:349–63.

9. LeVeque RJ, Mitchell IM, Stodden V. Reproducible Research for Scientific Computing: Tools and Strategies for Changing the Culture. Computing in Science & Engineering. 2012;14(4):13–7.

10. Afgan E, Baker D, Batut B, van den Beek M, Bouvier D, Čech M, et al. The Galaxy platform for accessible, reproducible and collaborative biomedical analyses: 2018 update. Nucleic Acids Research. 2018 Jul 2;46(W1):W537–44.

11. Bellec P, Lavoie-Courchesne S, Dickinson P, Lerch JP, Zijdenbos AP, Evans AC. The pipeline system for Octave and Matlab (PSOM): a lightweight scripting framework and execution engine for scientific workflows. Front Neuroinform [Internet]. 2012 [cited 2020 Jan 8];6. Available from: http://journal.frontiersin.org/article/10.3389/fninf.2012.00007/abstract

12. Gorgolewski K, Burns CD, Madison C, Clark D, Halchenko YO, Waskom ML, et al. Nipype: A Flexible, Lightweight and Extensible Neuroimaging Data Processing Framework in Python. Front Neuroinform [Internet]. 2011 [cited 2020 Jan 9];5. Available from: http://journal.frontiersin.org/article/10.3389/fninf.2011.00013/abstract

13. Oinn T, Addis M, Ferris J, Marvin D, Senger M, Greenwood M, et al. Taverna: a tool for the composition and enactment of bioinformatics workflows. Bioinformatics. 2004 Nov 22;20(17):3045–54.

14. Pestilli F, Hayashi S, Caron B, Vinci-Booher S. Brainlife. 2017.

15. Rex DE, Ma JQ, Toga AW. The LONI Pipeline Processing Environment. NeuroImage. 2003 Jul;19(3):1033–48.

16. The FIL Methods Group. SPM12 Manual. 2020.

17. Defaix F, Doyle M, Wetmore R. Version Control System for Software Development. Waterloo, ON; US 7,680,932 B2, 2010. p. 17.

18. Gorgolewski KJ, Auer T, Calhoun VD, Craddock RC, Das S, Duff EP, et al. The brain imaging data structure, a format for organizing and describing outputs of neuroimaging experiments. Sci Data. 2016 Dec;3(1):160044.

19. Haikin JS. Version control system for software code [Internet]. Fremont, CA; US6757893B1, 2004 [cited 2020 Jan 6]. Available from: https://patentimages.storage.googleapis.com/d0/4e/da/739afdc74e1bc0/US6757893.pdf

20. Spinellis D. Version Control Systems. IEEE Softw. 2005 Sep;22(5):108–9.

21. Ram K. Git can facilitate greater reproducibility and increased transparency in science. Source Code Biol Med. 2013 Dec;8(1):7.

22. Brown NCC, Wilson G. Ten quick tips for teaching programming. Ouellette F, editor. PLoS Comput Biol. 2018 Apr 5;14(4):e1006023.

23. Dudley JT, Butte AJ. A Quick Guide for Developing Effective Bioinformatics Programming Skills. PLoS Computational Biology. 2009;5(12):7.

24. Andersen LM. Group Analysis in FieldTrip of Time-Frequency Responses: A Pipeline for Reproducibility at Every Step of Processing, Going From Individual Sensor Space Representations to an Across-Group Source Space Representation. Front Neurosci. 2018 May 1;12:261.

25. van Vliet M. Seven quick tips for analysis scripts in neuroimaging. Markel S, editor. PLoS Comput Biol. 2020 Mar 26;16(3):e1007358.

26. Van Noorden R. The Trouble with Retractions. Nature. 2011 Jun 10;478(7367):26–8.

27. Niso G, Gorgolewski KJ, Bock E, Brooks TL, Flandin G, Gramfort A, et al. MEG-BIDS, the brain imaging data structure extended to magnetoencephalography. Sci Data. 2018 Dec;5(1):180110.

28. Pernet CR, Appelhoff S, Gorgolewski KJ, Flandin G, Phillips C, Delorme A, et al. EEG-BIDS, an extension to the brain imaging data structure for electroencephalography. Sci Data. 2019 Dec;6(1):103.

29. Clyburne-Sherin A, Fei X, Green SA. Computational Reproducibility via Containers in Psychology. MP [Internet]. 2019 Nov 12 [cited 2020 Feb 14];3. Available from: https://open.lnu.se/index.php/metapsychology/article/view/892

30. MATLAB. Natick, Massachusetts: The Mathworks Inc.; 2020.

31. Knuth DE. Literate Programming. Center for the Study of Language and Information; 1992.

32. Live Scripts and Functions [Internet]. MathWorks; 2020 [cited 2020 Dec 2]. Available from: https://uk.mathworks.com/help/matlab/live-scripts-and-functions.html

33. Potse M. matlabweb [Internet]. CTAN; [cited 2020 Dec 2]. Available from: https://ctan.org/pkg/matlabweb?lang=en

34. Jupyter Notebook [Internet]. Jupyter; 2020 [cited 2020 Dec 2]. Available from: https://jupyter.org/

35. Allaire J, Xie Y, McPherson J, Luraschi J, Ushey K, Atkins A, et al. rmarkdown: Dynamic Documents for R. [Internet]. R Studio; 2020 [cited 2020 Dec 2]. Available from: https://github.com/rstudio/rmarkdown

36. Xie Y. knitr: A General-Purpose Package for Dynamic Report Generation in R [Internet]. 2020. Available from: https://yihui.org/knitr/

37. Kery MB, Radensky M, Arya M, John BE, Myers BA. The Story in the Notebook: Exploratory Data Science using a Literate Programming Tool. In: Proceedings of the 2018 CHI Conference on Human Factors in Computing Systems - CHI ‘18 [Internet]. Montreal QC, Canada: ACM Press; 2018 [cited 2020 Jan 9]. p. 1–11. Available from: http://dl.acm.org/citation.cfm?doid=3173574.3173748

38. Oostenveld R, Fries P, Maris E, Schoffelen J-M. FieldTrip: Open Source Software for Advanced Analysis of MEG, EEG, and Invasive Electrophysiological Data. Computational Intelligence and Neuroscience. 2011;2011:1–9.

39. Oostenveld R. Wakeman-and-Henson-2015 [Internet]. Github; 2016. Available from: https://github.com/robertoostenveld/Wakeman-and-Henson-2015

